# Characterizing the human methylome across the life course: findings from eight UK-based studies

**DOI:** 10.1101/2021.09.18.460916

**Authors:** Esther Walton, Riccardo Marioni, Hannah R Elliott, Simon R Cox, Ian J Deary, Alun D Hughes, Therese Tillin, Meena Kumari, Tom Woofenden, Juan E Castillo-Fernandez, Jordana T Bell, Alissa Goodman, George Ploubidis, Kate Tilling, Matthew Suderman, Tom R Gaunt, Erin C Dunn, Andrew Smith, Caroline L Relton

**Author notes:** corresponding author: Dr Esther Walton, Department of Psychology, University of Bath, Bath, BA2 7AY, United Kingdom. shared authorship (Andrew Smith, Caroline L Relton).

## Abstract

Variation in DNA methylation (DNAm) is associated with multiple biological processes that track growth and development, ageing and age-related diseases. However, there is little understanding of what constitutes typical patterns of DNAm variation and how these patterns change across the life course. In this study, we synthesised a map of the human methylome across the life course, focussing on changes in variability and mean DNAm.

Harmonizing DNAm datasets across eight longitudinal and cross-sectional UK-based studies, we meta-analysed n=13,215 blood samples from n=7,037 unique individuals from birth to 98 years of age. Changes in CpG-specific variability and means were described across the life course using a meta-regression framework. CpG-specific associations of variability or mean DNAm in relation to the likelihood of association with 100 traits linked to environmental exposures, health and disease were tested within and across ten developmental age bins across the life course.

Age was linked to DNAm variability at 29,212 CpG sites. On average, we observed a 1.26 fold increase in DNAm variability per year across the life course. 33,730 CpGs displayed changes in mean DNAm, with 64% of these loci showing decreases in DNAm over time. CpG sites linked to traits were in general more variable across the life course.

Our study provides, for the first time, a map of the human methylome across the life course, which is publicly accessible through a searchable online database. This resource allows researchers to query CpG-specific trajectories from birth to old age and link these to health and disease.

## Introduction

A wealth of research has emphasized the importance of epigenetic mechanisms, such as DNA methylation (DNAm), in both developmental processes and a range of diseases, including cancer and neuropsychiatric outcomes (Baglietto et al., 2017; Barker et al., 2018; Smith and Meissner, 2013). Perhaps surprisingly, DNAm displays a strong association with aging to such a degree that we are now able to derive reliable and accurate predictors of chronological age in children and adults using DNAm alone (Hannum et al., 2013; Horvath, 2013; Ryan et al., 2020). However, there is still no clear understanding of what constitutes normative or healthy (versus atypical or disease-linked) patterns of DNAm variation and change across the life course.

There is substantial evidence for both intra- and inter-individual CpG-specific variability and change over time in healthy individuals (Milnik et al., 2016; Oh and Petronis, 2021; Wang et al., 2012). Currently, the largest study focussing on methylation trajectories from birth to late adolescence, Mulder et al. (2021) pooled data from two birth cohorts with a combined sample size of 2,348 participants and identified widespread and often non-linear changes in DNAm throughout childhood. Investigating CpG-specific associations with age in adulthood, McCartney et al. utilised data from the Generation Scotland study, including a discovery sample of 2,586 unrelated individuals and a replication sample of 4,450 individuals aged 18 to 93 (2019). Using this cross-sectional, adult-only sample, researchers identified age associations in up to 31% of the CpGs analysed, where age was found to associate with some measurable degree of DNAm variability. Notably, previous studies were either cross-sectional, or focussed predominantly on age associations with changes in *mean* levels of methylation. Restricted by the age range of the underlying study sample, they also characterised methylation over a more restricted time period.

We aimed to build a map of the human methylome across the whole life course (birth to 100 years of age) by leveraging data from multiple longitudinal and cross-sectional UK-based studies. To do so, we developed harmonization protocols to ensure comparability of methylation data across studies, allowing us to derive meaningful conclusions about the life course methylome. We characterize this map in terms of changes in variability as well as mean changes across the life course. Results are fully searchable through our online resource MATL (‘Methylation across the life course’) at http://browser.ariesepigenomics.org.uk/. With this tool, we enable researchers to query CpG-specific methylation patterns from birth to old age to 1) assess the dynamic nature of the human methylome at specific developmental periods and across the life course; 2) narrow their own analysis search space to CpG sites that show changes in variability over the life course; and 3) link these patterns to traits of health and disease.

## Methods

### Sample Descriptions

We pooled ten datasets from eight birth or population-based UK-based cohort studies (Table 1 and SM section 1.1), for a total of n=13,215 blood-based DNAm samples from n=7,037 unique individuals. Age ranged from birth to 98 years.

**Table 1.**
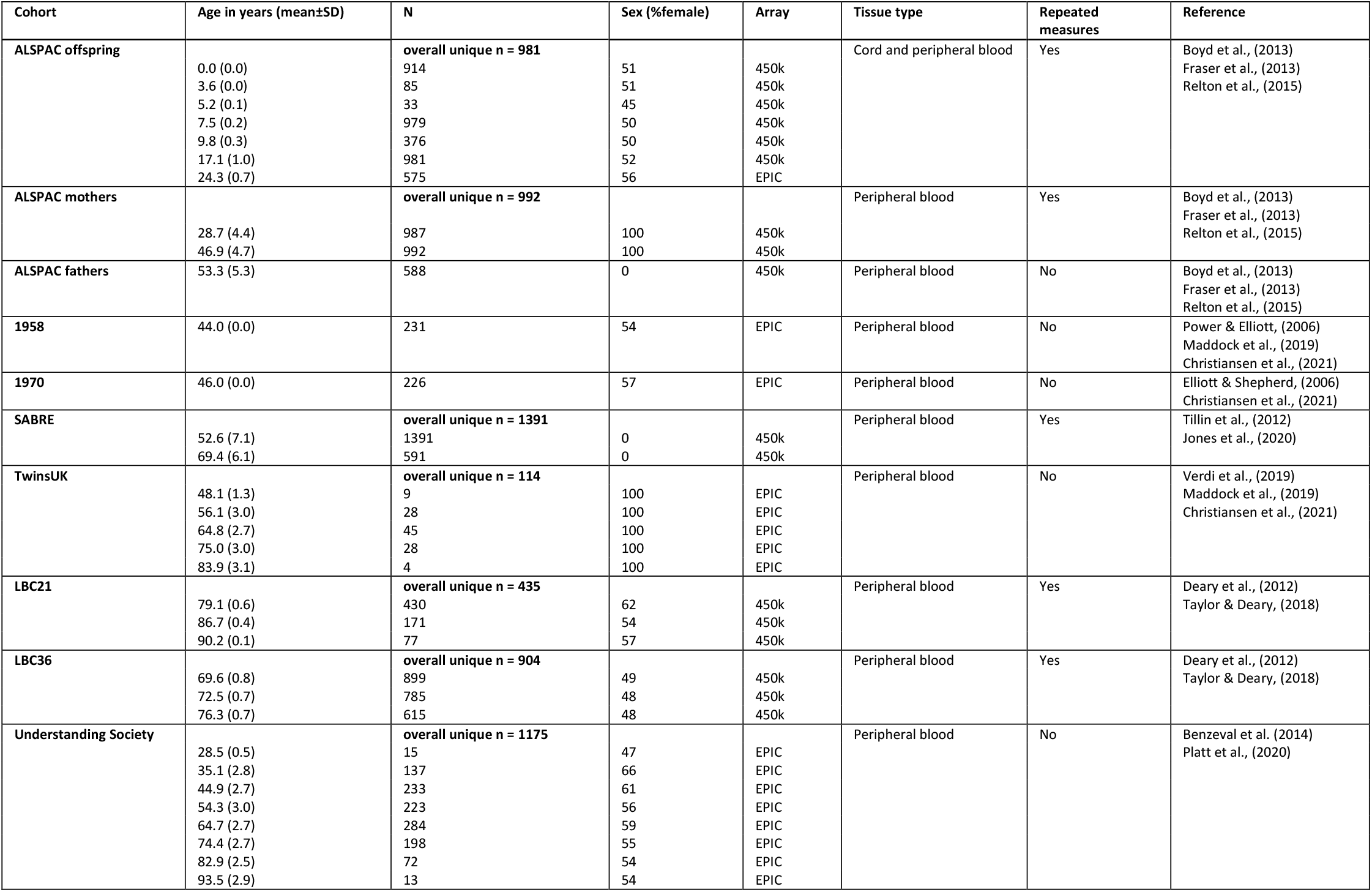
Sample descriptives of the eight UK-based cohort studies, split into 33 datasets for the current study.

Five of these eight cohorts were longitudinal in design with at least two repeated measures of DNAm available (range: 2 to 7 time points of DNAm). Four studies included females or males only; the remaining studies contained a roughly equal ratio of males:females. DNAm data were generated using Illumina EPIC or 450k HumanMethylation BeadChip arrays.

The two cross-sectional, non-birth cohort studies (TwinsUK and *Understanding Society*) spanned several age decades and were therefore split into ten-year age bins, resulting in a total of 33 datasets used for analysis in the current study (Table 1).

### DNA methylation data preprocessing and harmonization

We used *meffil* (Min et al., 2018), an R package designed for efficient quality control, normalization and epigenome-wide association studies of large samples of Illumina Methylation BeadChip microarrays to pre-process methylation data for nine datasets. *Understanding Society* data was already pre-processed externally (see SM section 1.1). By harmonizing pre-processing we were able to reduce heterogeneity across cohorts due to diverse pre-processing pipelines (SM section 1.4).

Importantly, *meffil* allows users to normalize datasets together using an approach called functional normalization without needing to share individual-level data. The *meffil* workflow divides functional normalization into three steps: 1) creation of raw data summaries and exclusion of DNAm samples failing quality control within each dataset; 2) pooling and normalization of the data summaries across all datasets and 3) normalization of each dataset using normalized summaries and exclusion of CpG sites failing quality control. Functional normalization preserves mean and variance structure of the data. For further details, see SM section 1.2. For numbers of samples and CpG sites excluded from each cohort, see SM Table 1. Detailed scripts are available at https://github.com/stegosaurusrox/Lifecourse_Methylome.

### Statistical analysis

For a flow chart of our analysis protocol, see SM Figure 2. We extracted seven summary statistics per CpG across all individuals in a given dataset, using the *quantile* and the *stats*.*desc* functions in the pastecs R package: minimum, maximum, median, mean, standard deviation, 25th and 75th percentiles.

Our main aim was to describe CpG-specific changes in variability and mean tendency across the life course and therefore, we focussed on the following two summary statistics in particular - standard deviation (SD) and mean. Users can visualize these two characteristics using MATL (‘Methylation across the life course’), the developed online tool available under http://browser.ariesepigenomics.org.uk/. Users are also provided with a table listing all seven descriptive statistics including the median and interquartile range.

### Change in variability and mean DNA methylation over the life course

To assess trends in the variability of DNAm over the life course (indexed by the standard deviation; SD), we conducted meta-regressions of SD DNAm (outcome) on age (predictor) for each CpG site using summary statistics. We used a multi-level meta-regression model to account for two potential sources of correlation within measures of DNAm at the same CpG site. The first source considered was correlation *within* a cohort, due to the presence of repeated measures on the same individuals within a cohort. The second source considered was correlation *between* cohorts within the same study (e.g. mothers, fathers and offspring within ALSPAC) due to i) the possibility of within-study similarities in pre-processing that was not fully removed during harmonization, and ii) inter-generational relationships between individuals within the same study. Full details of the model are given in Supplementary Material section 1.5.

The multi-level model is specified such that the fixed effects, random effects, and error terms can be either positive or negative. On the other hand, the dependent variable measures, the standard deviations, are always positive. We applied a log transformation to the standard deviations, so that both sides of the model could match. Throughout this study, we refer to the log transformed SD as SD_DNAm_ for simplicity. In this model, we controlled for effects of mean methylation levels due to the assumption that the variance is influenced by the mean (but not vice versa).

We also carried out meta-regressions of mean DNAm on age for each CpG site using summary statistics. In this model, the dependent variable measures, the mean methylation levels, are a proportion. We applied a logit transformation to mean methylation levels so that both sides of the model could match. For full details, see SM section 1.5.

In our main analyses, we did not control for cell type composition, as cell type changes have been shown to be a meaningful component of the ageing process (Valiathan et al., 2016) and correcting for cell type might remove these ageing effect, which we sought to explicitly model here. However, we also report cell-type adjusted results throughout the results and in the supplementary materials as a sensitivity analysis.

To test for sex-specific associations, we fitted separate random-effects models in males and females. To ascertain sex-interaction effects, we combined the two single-sex subsets using a fixed-effects model, because the (residual) heterogeneity within each subset has already been accounted for by fitting random-effects models above. Fitting separate single-sex models also allowed the amount of heterogeneity within each set to be different (Rubio-Aparicio et al., 2020). The Y chromosome was excluded from these analyses.

Forty-nine percent of all assessed CpG sites (n=899,096) were present in all 33 cohorts (as well as in all 30 all-female and 26 all-male cohorts), with a further 45% present in all 16 cohorts (as well as in all 16 all-female and 11 all-male cohorts; SM Table 4). Epigenome-wide levels of significance were defined as p < 1*10-7.

Gene ontology analyses were carried out using the gometh function in the missMethyl R package (version 1.20.4), which takes into account the probability of significant differential methylation due to numbers of probes per gene.

Enrichment for methylation quantitative trait loci (mQTLs) was tested using mQTL summary data available for 420,509 CpGs through the GoDMC database (http://mqtldb.godmc.org.uk/). Clumped cis mQTLs were defined based on a threshold of p<1*10^−8^, while trans associations were based on p<1*10^−14^. The observed counts of age-associated CpGs with mQTLs (versus all age-associated CpGs) was contrasted to the expected counts, based on all mQTL-linked CpGs (n=190,102) versus all tested CpGs in GoDMC.

### Age-specific enrichment of trait-associated CpGs

To investigate if CpG sites, which have been linked to environmental exposures, health and disease traits on an epigenome-wide level, have an age-specific profile compared to un-associated CpGs, we downloaded the EWAS catalogue (http://www.ewascatalog.org/; accessed Jul 19, 2019) and selected probes that have been linked to at least one trait at an epigenome-wide level (p < 1*10^− 7^) in blood tissue (referred to as CpG_EWAS_). We then grouped these CpGs into 12 time bins (birth, 0.1-5 years, 6-10 years and then every decade), according to the mean age of the study that reported the finding. No associations were listed for any age group +80y, leaving 10 time bins for analysis. We identified 128 unique EWAS with a sample size of at least 100 individuals, reporting 103,456 unique associations for 100 traits. There were more publications focussing on birth and on samples between the age of 40-70 years than on any other age period. Studies on these age periods were also the largest in sample size and reported the greatest number of associations (SM Figure 11 and SM Table 5). As all listed studies were based on the 450k array, our analyses were restricted to probes on this array only.

## Results

### Widespread increase in CpG-specific variability across the life course

Our main focus was to investigate how SD_DNAm_, a measure of CpG-specific variability, changed across the life course. An epigenome-wide meta-regression of SD_DNAm_ on age revealed wide-spread age associations across the genome. Specifically, 29,212 CpGs (3.25% of all tested probes) passed a significance threshold of 1*10^−7^ (lambda = 3.02; Figure 1A, SM Figure 6A and SM Table 6). The majority of these associations (98.74%) were positive, indicating an increase in CpG-specific variability over the life course (Figure 1A and 1B). On average, we observed a 1.26 fold increase in SD_DNAm_ per year over and above any increase expected from concurrent changes in mean methylation. This average effect estimate decreased to 1.23 once controlling for cellular composition (SM section 2.1).

**Figure 1.**
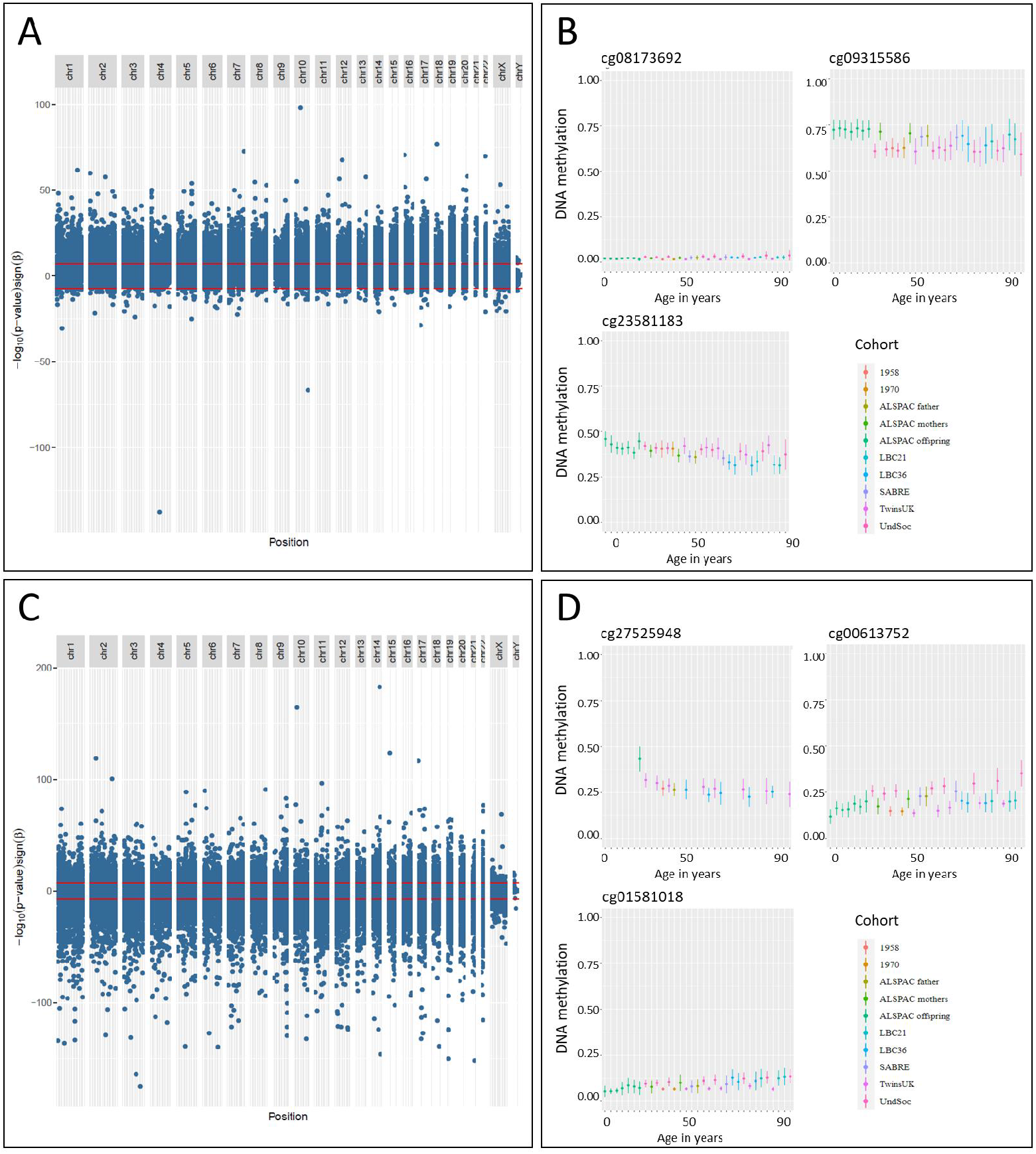
On the left, Miami plots of age associations with A) SD_DNAm_ levels and C) mean_DNAm_ levels across the lifecourse with chromosome position on the x-axis and -log(p-value)*sign(β) on the y-axis. On the right, mean methylation (±SD) of selected CpG sites showing changes in B) SD_DNAm_ levels or D) mean_DNAm_ levels across the lifecourse.

The largest increase in variability across all 33 datasets, after correcting for changes in mean methylation, was observed for cg08173692 (b_log(SD)_= 0.64; 95% CI: [0.47; 0.80] corresponding to a 1.89 fold increase per year; Figure 1B). This CpG is located proximal to the Proline-rich Protein gene *PRR18*. We also found evidence for a functional link between cg08173692 and *PRR18*. Thirty-three of the 46 mQTLs for cg08173692 listed in the GoDMC database were also eQTLs for *PRR18* (SM Table 7). The association between age and SD_DNAm_ was slightly stronger for overall more variable CpG sites (rho=0.32, SM Figure 8A). In other words, higher overall variability related to increased variability over time.

Age cohort effects did not explain all variance in SD_DNAm_ with residual heterogeneity remaining for 86.55% of probes overall (mean tau^2^=0.11) and for 67.51% of those associated with age (mean tau^2^=0.05). Residual heterogeneity was largely reduced once controlling for cell type (mean tau^2^=0.008). Gene ontology analysis indicated processes such as synaptic signalling to be linked to changes in SD_DNAm_ (SM Table 8). SD_DNAm_-associated CpGs were not enriched for or depleted of mQTLs (*χ*^2^=0.35, *p*=0.55), suggesting statistically indiscernible genetic contribution to age-related methylation variability.

### Widespread changes in mean methylation levels across the life course

Age was associated with mean_DNAm_ in n=33,730 CpG sites (3.75% of all probes; Figure 1C, SM Table 9). As expected, associations were widespread with high genomic inflation (lambda = 3.41) (SM Figure 6B). Mean methylation changes in probes were largest in infancy, changing on average by 3% percentage points per year compared to 0.1% in adulthood. See Figure 1D for examples of CpGs sites with age associations on mean_DNAm_. Mean_DNAm_ decreased with age in 64.33% of these probes. The effect estimate of the association between age and mean_DNAm_ was very weakly (yet negatively) correlated with mean_DNAm_ (rho=-0.22; SM Figure 8B), possibly representing floor or ceiling effects. That is, age associated with mean_DNAm_ slightly more positively (or less negatively) for less methylated CpG sites, but slightly more negatively (or less positively) for highly methylated CpG sites. Residual heterogeneity between studies was present for almost all age-associated CpGs (99.03%, mean tau^2^ = 0.06) but slightly lower compared to all probes on the array (mean tau^2^ = 0.10). Additionally correcting for cellular composition further decreased these heterogeneity levels (mean tau^2^=0.005; SM section 2.1 and SM Table 9). Gene ontology analyses highlighted processes linked to nervous system development (SM Table 8). Mean_DNAm_-associated CpGs were depleted of mQTLs (*χ*^2^=710.9, p=1.28*10^−156^), suggesting that age-associated CpG sites are under less genetic control than CpG sites overall. Overall, these results indicate widespread, but small changes in mean methylation across the life course, with a trend towards a decrease in methylation with age, rather than an increase.

### Age associations with mean methylation confirm previous findings

The proportion of CpGs associated with age was much smaller than reported elsewhere (e.g. 3.75% versus 18.96% in McCartney et al. (2019) and 51.6% in Mulder et al. (2021)). We therefore undertook a more detailed comparison with associations reported in one of these previous studies, which were obtained from a large, non-overlapping sample of adults from GenerationScotland (2019).

Regression coefficients for our 33,730 age-associated CpG sites correlated very highly between studies (replication.1: rho=0.93; replication.2: rho=0.92; SM Section 2.2). The reverse was also true: regression coefficients of age-associated CpG sites reported in McCartney et al. (2019) correlated strongly between studies (replication.1: rho=0.88; replication.2: rho=0.87). The strength of these correlations decreased slightly once controlling for cellular composition in our analysis (rho’s between 0.74 to 0.81; see SM section 2.2 and SM Figure 7). Furthermore, the proportion of probes with decreasing mean_DNAm_ over time was largely consistent across studies (64.33% versus 55.66% in McCartney et al. (2019)).

These findings suggest our identified CpGs reflect a subset of true positives, though we were possibly slightly underpowered to detect small associations due to a meta-regression approach based on summary data from multiple studies compared to an individual-level analysis performed in a single study by McCartney et al (2019).

### Age associations are independent of probe-specific reliability

Our meta-analysis results were based on data measured on two different microarrays (450k and EPIC), previously shown to display a mean inter-array reliability as low as 0.21 across CpG sites (Sugden et al., 2020). We therefore assessed – using reliability measures as reported in Sugden et al. (2020) – the degree to which our findings could be influenced by low probe-specific reliability. *Within* each of the 33 cohorts, we could replicate previous findings, namely 1) a positive association between the SD (untransformed) and reliability and 2) an inverse-U shaped relationship between mean methylation (untransformed) and reliability (SM Figure 9A and 9B). However, our meta-regression results of SD_DNAm_ and mean_DNAm_ on age *across* cohorts were independent of reliability estimates (SM Figure 9C and 9D), indicating that our results were not driven by low inter-array reliability.

### Little evidence for sex-specific effects on DNAm levels and variability

Separate analysis in females revealed 15,385 CpGs (versus 15,259 in males) linked to SD_DNAm_ with a moderate degree of overlap (SM Figure 10A). Age-by-sex-interaction effects were only detected in 112 CpG sites. Roughly a third of these were located on the X chromosome, where the association between age and SD_DNAm_ was generally weaker in males (mean beta_males_=-0.29 corresponding to a 25% relative decrease in the age effect in males). Across the autosomes, associations were overall larger in males (mean beta_males_=0.56, corresponding to a 75% relative increase in the age effect in males compared to females; SM Table 10).

For mean_DNAm_, we discovered slightly more age-associated CpGs in males (n_CpG_=31,537) than in females (n_CpG_=19,963; SM Figure 10B). Testing formally for an age-by-sex-interaction effect identified 1,279 CpG sites, of which 81.86% displayed a more positive (i.e. sharper increase in mean level over the life course) age association in males compared to females (SM Table 11). We did not observe the same clustering of effects on the X chromosome, as we did for SD_DNAm_.

Together, these findings suggest few sex-specific associations between age and SD_DNAm_, which were mainly localized to the X chromosome. Sex-specific age associations with mean_DNAm_ were more widespread across chromosomes, but still somewhat rare.

### Age-specific enrichment of trait-associated CpGs

As a final step, we investigated if CpG sites associated with traits have different levels of SD_DNAm_ and mean_DNAm_ than non-trait associated CpGs. We similarly asked if age associations with SD_DNAm_ or mean_DNAm_ differed between trait- and non-trait associated CpGs.

First, we found that trait-associated CpG sites tended to have higher SD_DNAm_ at most ages except in childhood and middle age (50s; Figure 3A and SM Table 12). In other words, trait-associated CpG sites were more variable, but not consistently so across all ages.

**Figure 3.**
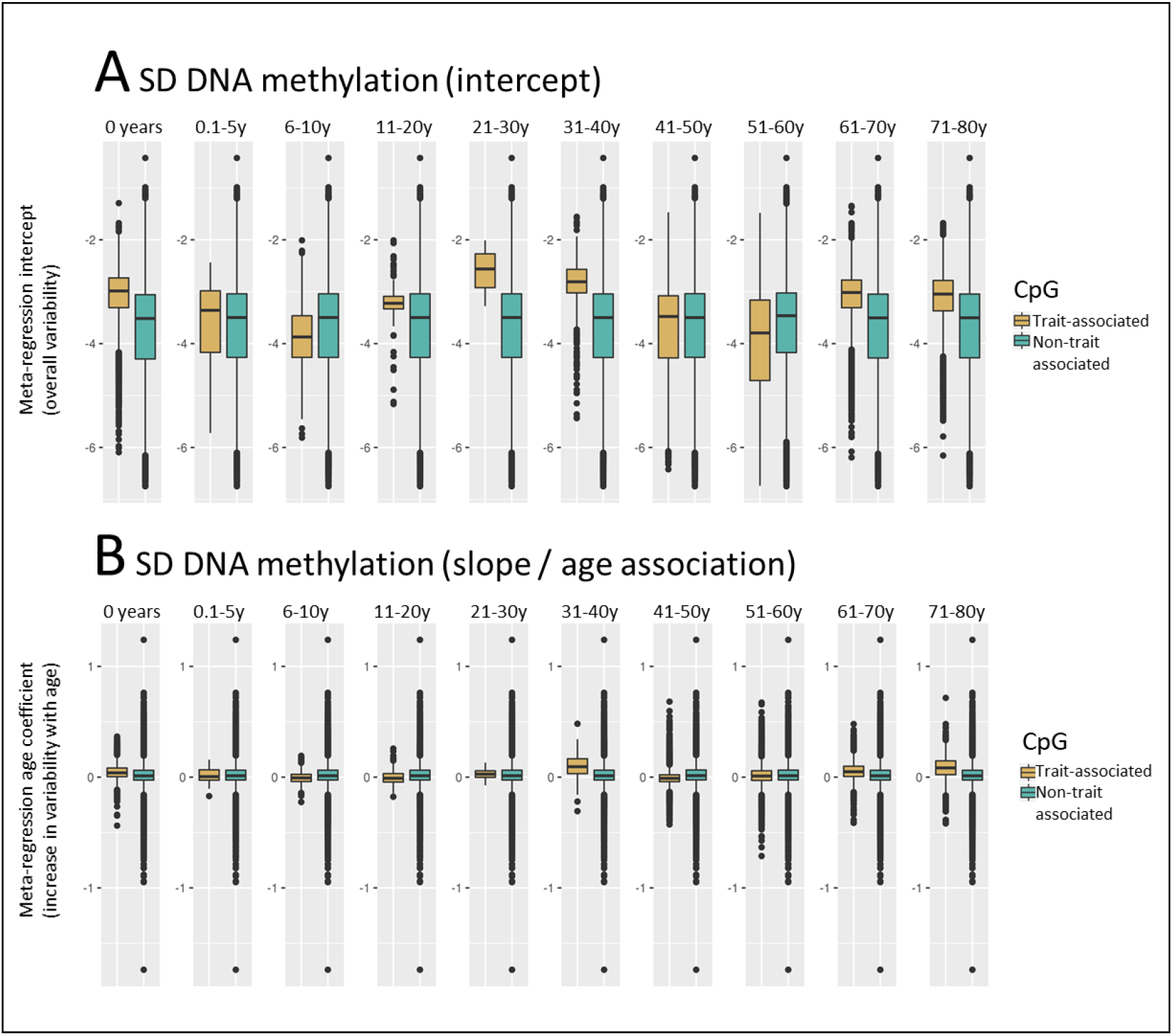
A) SD_DNAm_ (intercept) and B) the change in SD_DNAm_ (slope) across the life course in CpGs reported in EWAS versus un-reported CpGs by age period (in panels). Values on the y-axis refer to a regression with log(SD) as an outcome. Larger values indicate an increase in variability. Trait-associated CpGs were identified using the EWAS catalog database (http://ewascatalog.org/).

The patterns for all other associations were more complex. Although trait-associated CpGs were generally more variable, they showed both further increases, but also decreases in variability over the life course compared to unlinked CpGs. The same was true for mean_DNAm_ levels (Figure 3B, SM Figure 12A and 12B and SM Table 12).

### Lifecourse Methylome Browser

We made all summary results openly available through our online resource MATL at http://browser.ariesepigenomics.org.uk/. Researchers can query CpG-specific trajectories from birth to old age, for example to narrow their own analysis search space to CpG sites that show changes in variability over the life course.

## Discussion

### Summary of findings

The main finding of this study is that age-linked changes in DNA methylation variability and mean levels occur across the methylome. DNA methylation variation changes with age at 3.25% of CpG sites, almost always increasing with age. DNA methylation levels change across the life-course at 3.75% of CpG sites, two-thirds of these decreasing with age. Genetic influences shaped age-related mean methylation levels, but not methylation variability. Sites with changing variation were more likely to have been linked to traits than those with more constant variation. However, there was a large degree of residual heterogeneity between studies suggesting that individual CpG sites are characterised by unique patterns of variability across the life course. Our study provides, for the first time, a map of the human methylome across the life course, which is publicly accessible through a searchable online database at http://browser.ariesepigenomics.org.uk/. With this resource, researchers can query CpG-specific trajectories from birth to old age, link these to health and disease traits and assess the dynamic nature of the human methylome at specific developmental periods and across the life course.

### Age associations with SD DNA methylation

We found that the vast majority of probes showed a positive association between age and SD of DNA methylation, indicating an increase in CpG-specific variability across the lifespan, in line with previous studies (Slieker et al., 2016; Talens et al., 2012). This variability could have important downstream transcriptional effects, as a recent study reported that promoters with increased methylation heterogeneity related to increased transcriptional heterogeneity of their associated genes (Hernando-Herraez et al., 2019).

Findings from two studies suggest that the increase in DNA methylation variability with age might be driven by an increase of environmental or stochastic influences over time, rather than a decrease in stable genetic influences. Gaunt et al. (2016) showed that the average SNP heritability decreased only slightly from 24% in childhood to 21% in middle age; and Talens et al. (2012) reported that methylation variability increased with age in 230 MZ twin pairs (18-89 years), further suggesting that environmental (or stochastic) factors might contribute to these phenomena. It is possible that our findings of increased variability relate to epigenetic drift, which is often considered a hallmark of ageing and thought to be driven by the loss in fidelity of epigenetic marks over time (Mendelsohn and Larrick, 2017). Future studies should assess the degree to which increases in variability (and epigenetic drift) are a contributor to biological ageing or merely a bystander phenomenon.

The largest increase in variability in a single CpG site was observed at cg08173692 proximal and functionally linked to the Proline-rich Protein 18 *(PRR18)* gene. Interestingly, this gene is linked to genetic variation associated with human longevity (Yashin et al., 2010).

Similarly, a study of age-by-sex-specific variability in DNA methylation found marked sex differences in another Proline-rich Protein gene coding for PRR4 which diverged further with age (Yusipov et al., 2020). Proline is known to pay an important role in the immune response, and proline supplementation has shown to increase life expectancy in microbial populations (Canfield and Bradshaw, 2019). The proline-rich Akt substrate is the inhibitory subunit of the mTOR complex 1 (mTORC1). The inhibition of this complex is currently the only mechanism known for increasing lifespan in all model organisms studied (Saxton and Sabatini, 2017). Future studies are needed to further investigate the role of DNA methylation as an aging mechanism that regulates proline-rich proteins in the mTOR signalling hub.

### Age associations with mean DNA methylation

Over the lifespan, mean methylation of a considerable proportion of probes changed with age. Slightly more CpG sites showed a decrease in methylation than an increase, consistent with previous research (McCartney et al., 2019; Mulder et al., 2021). However, the overall proportion of probes we identified was much smaller than in these previous studies (3.75% versus 20% and 51.6% respectively). It is possible that our meta-regression approach was underpowered to detect small associations compared to an individual-level analysis applied in McCartney et al. (2019) and Mulder et al. (2021). However, it is also possible that CpG-specific changes are more developmentally specific. CpGs that fluctuate in mean methylation during childhood might not continue to do so in adulthood and vice versa. In fact, recent research suggests that age-related CpG-specific changes are logarithmic, with steep changes in childhood that taper off in adulthood (Snir et al., 2019). As previous research was confined to specific stages in the life course, either the period from birth to adolescence (Mulder et al., 2021) or adulthood only, (McCartney et al., 2019), these studies might have been better able to detect these developmentally specific changes, which our models did not explicitly examine. Together with the large degree of heterogeneity, reported in the current study, this suggests that the human methylome might not change linearly throughout the life course, further emphasizing the need to map methylomic patterns with more granularity.

### Sex-specificity

We found weak age-by-sex interaction effects on mean DNA methylation or variability across the life course, which partially aligns with previous findings. The Copenhagen puberty cohort found that 457 methylation sites – similar to the number we report – are tightly associated with pubertal transition and altered hormone levels, with stronger associations in boys than in girls (Almstrup et al., 2016). However, Han et al. (2019) reported age-by-sex interaction in around 13.17% of probes during puberty, which is a much higher estimate than what we found. It might be that these changes are specific to puberty or narrow during later developmental stages such as menopause and hence do not remain detectable across the life course. A study analysing methylation data of 703 newborns (one of the datasets used in the current work) also reported weak sex effects on variability in only 240 CpGs (Staley et al., 2020), further highlighting the need for a life course approach to study the dynamic nature of the human DNA methylation.

### Developmental enrichment of trait-associated CpGs

It was evident that CpGs, which have already been linked to traits, were consistently more variable in most developmental stages throughout the life course compared to CpG sites that have not yet been linked to phenotypes. This aligns with Sugden et al. (2020), who showed that CpG sites with increased reliability (which itself was correlated with methylation variance) were more consistently associated with smoking across several studies. Our study extended that finding by showing that the link between increased variance and trait-associability might hold for traits other than smoking and across the life course. In contrast, however, we observed some independence from CpG reliability. It is also possible that our finding represents a statistical by-product (or ‘survival’ effect) as only CpGs that show discernible inter-individual variation would be detected as having an association with a particular trait, while largely invariant probes would go undetected.

We were unable to see the same consistent associations with mean methylation levels, suggesting that studying variability – rather than mean differences in methylation – might be an additional aspect that could be considered when assessing the degree to which DNA methylation links to traits across the life course.

### Overlapping trends

We observed roughly similar numbers of CpG sites with age-associated DNA methylation levels or variance. These two sets of CpG sites were largely non-overlapping, which is partially due to design as age association with variability were controlled for concurrent changes in mean methylation. This is only partially in line with reports by Slieker et al. (2016), who reported i) four times as many probes linked to mean levels than to variability and ii) a 48.2% overlap between CpGs sites that were age-related variably methylated and at the same time showed a change in mean methylation levels with age. In their study, the authors covaried for cell type and sex effects, which might account for the observed differences and suggests that some of the age effects on variability, but not on mean methylation, might relate to changes in cell type composition or sex.

### Limitations

Our findings should be considered in light of the following limitations. The genomic inflation factor, the deviation in the distribution of the observed and expected test statistic, was shown to be elevated. In this study, this is probably explained by a poly-epigenic signal that is shared across many CpG across the genome, in line with previous reports (McCartney et al., 2019). However, high genomic inflation factors can also be caused by population stratification or strong associations between CpG sites.

In our main analysis, we did not control for cellular composition, as changes in cell type are part of the aging process (Valiathan et al., 2016). In line with this, we observed an attenuation of effect estimates (especially on mean methylation) and a reduction in residual between-cohort heterogeneity, when controlling for cell type. However, the overall patterns of increased variability and more widespread decreases than increases in mean methylation across the life course remained similar. Yuan et al. (2015) - using data from 656 samples from individuals aged 19 to 101y – also demonstrated that the age-associated increase in variability in methylation is largely independent of the DNA methylation variations found between blood cell types with 69% of methylation sites remaining stably associated with age, after cell type adjustment. Even higher concordance rates of 78.4% were observed comparing sets of variably methylated probes, obtained from whole blood vs purified monocytes (Slieker et al., 2016). Furthermore, our uncorrected results correlated strongly with those of McCartney et al. (2019), which were controlled for cell type.

Lastly, our analyses were based on pooling data from two different microarrays and previous studies have shown low inter-array reliability of methylation measurement (Bose et al., 2014; Dugué et al., 2016; Sugden et al., 2020). We developed a rigorous harmonization protocol to minimize these effects and subsequently found little evidence that our results were affected by low reliability, but this needs further validation as more and more studies across the age ranges are becoming available using the EPIC array.

## Conclusion

In summary, by drawing together harmonized DNA methylation data from eight longitudinal and cross-sectional UK-based studies with a total of n=13,215 samples from n=7,037 unique individuals aged 0 to 98, we mapped the human methylome across the life course and described widespread changes in variability and mean methylation across the life course, which were largely sex-independent. Trait-associated CpGs, identified from previously published EWAS studies, tended to be more variable. Our data have been curated as an openly accessible and searchable online database, allowing researchers to query CpG-specific trajectories from birth to old age, moving forward research in the field of life course epigenetics.

## Acknowledgements

ALSPAC

We are extremely grateful to all the families who took part in this study, the midwives for their help in recruiting them, and the whole ALSPAC team, which includes interviewers, computer and laboratory technicians, clerical workers, research scientists, volunteers, managers, receptionists and nurses.

TwinsUK

We would like to thank the twins for their participation.

1958 and 1970

The authors thank the 1958 and 1970 birth cohort study members.

Understanding Society

*Understanding Society* is an initiative funded by the Economic and Social Research Council and various Government Departments, with scientific leadership by the Institute for Social and Economic Research, University of Essex, and survey delivery by NatCen Social Research and Kantar Public. The research data are distributed by the UK Data Service. The epigenetics methylation data were analysed by the University of Exeter Medical School (MRC grant K013807) and further facilitated by the University of Essex School of Biological Sciences. Information on how to access the data can be found on the *Understanding Society* website https://www.understandingsociety.ac.uk/.

Lothian Birth Cohorts of 1921 and 1936

The authors thank all LBC1936 and LBC1921 study participants and research team members who have contributed, and continue to contribute, to ongoing LBC studies.

## Funding

This project is funded by CLOSER, whose mission is to maximise the use, value and impact of longitudinal studies. CLOSER was funded by the Economic and Social Research Council (ESRC) and the Medical Research Council (MRC) between 2012 and 2017. Its initial five-year grant was extended to March 2021 by the ESRC (grant reference: ES/K000357/1). The funders took no role in the design, execution, analysis or interpretation of the data or in the writing up of the findings. www.closer.ac.uk

E. Walton is also supported by EarlyCause, funded through the European Union’s Horizon 2020 research and innovation programme (grant nº 848158). Dr. Dunn is supported by the National Institute of Mental Health of the National Institutes of Health under Award Number R01MH113930. The content is solely the responsibility of the authors and does not necessarily represent the official views of the National Institutes of Health.

### ALSPAC

The UK Medical Research Council and Wellcome (Grant ref: 217065/Z/19/Z) and the University of Bristol provide core support for ALSPAC. This publication is the work of the authors and E.Walton will serve as guarantors for the contents of this paper. A comprehensive list of grants funding is available on the ALSPAC website (http://www.bristol.ac.uk/alspac/external/documents/grant-acknowledgements.pdf). This research was specifically funded by BBSRC (BBI025751/1 and BB/I025263/1), National Institute of Child and Human Development grant (R01HD068437), NIH (5RO1AI121226-02), CONTAMED EU collaborative Project (212502), LLHW via MRC (G1001357) and Wellcome Trust (WT092830/Z/10/Z). KT, MS, HRE, TG and CLR are members of the Integrative Epidemiology Unit which receives funding from the U.K. Medical Research Council and the University of Bristol (MC_UU_00011/3, MC_UU_00011/5).

### TwinsUK

TwinsUK was funded by the Wellcome Trust; European Community’s Seventh Framework Programme (FP7/2007-2013) and also receives support from the National Institute for Health Research (NIHR)-funded BioResource, Clinical Research Facility and Biomedical Research Centre based at Guy’s and St Thomas’ NHS Foundation Trust in partnership with King’s College London. The study received additional support from the ESRC (ES/N000404/1 to JTB).

### 1958 and 1970

These data were supported by the Economic and Social Research Council/Biotechnology and Biological Sciences Research Council (ES/N000404/1).

### SABRE

The SABRE study was funded at baseline by the Medical Research Council, Diabetes UK, and the British Heart Foundation. At follow up the study was funded by the Wellcome Trust (082464/Z/07/Z), the British Heart Foundation (SP/07/001/23603, PG/08/103, PG/12/29/29497 and CS/13/1/30327) and Diabetes UK (13/0004774). Methylation analysis in the SABRE cohort was supported by a Wellcome Trust Enhancement grant (082464/Z/07/C). ADH received support from the National Institute for Health Research University College London Hospitals Biomedical Research Centre. ADH and NC work in a unit that receives support from the UK Medical Research Council (MC_UU_12019/1). Support has also been provided at follow-up by the North and West London and Central and East London National Institute of Health Research Clinical Research Networks.

### Understanding Society

These data are from *Understanding Society*: The UK Household Longitudinal Study, which is led by the Institute for Social and Economic Research at the University of Essex and funded by the Economic and Social Research Council (Grant Number: ES/M008592/1). The data were collected by NatCen and the genome wide scan data were analysed by the Wellcome Trust Sanger Institute. Information on how to access the data can be found on the *Understanding Society* website https://www.understandingsociety.ac.uk/. Data governance was provided by the METADAC data access committee, funded by ESRC, Wellcome, and MRC. (2015-2018: Grant Number MR/N01104X/1 2018-2020: Grant Number ES/S008349/1).

### Lothian Birth Cohorts of 1921 and 1936

Information on the Lothian Birth Cohorts, including data access, can be found on the website: https://www.ed.ac.uk/lothian-birth-cohorts. LBC1921 was supported by the UK’s Biotechnology and Biological Sciences Research Council (BBSRC), a Royal Society–Wolfson Research Merit Award to I.J.D, and the Chief Scientist Office (CSO) of the Scottish Government’s Health Directorates. The LBC1936 is supported by Age UK (Disconnected Mind program), the Medical Research Council (G0701120, G1001245, MR/M013111/1, MR/R024065/1), and the University of Edinburgh. Methylation typing was supported by Centre for Cognitive Ageing and Cognitive Epidemiology (Pilot Fund award), Age UK, The Wellcome Trust Institutional Strategic Support Fund, The University of Edinburgh, and The University of Queensland. This work was conducted in the Centre for Cognitive Ageing and Cognitive Epidemiology, which is supported by the Medical Research Council and Biotechnology and Biological Sciences Research Council (MR/K026992/1), and which supported I.J.D. R.E.M. is supported by Alzheimer’s Research UK major project grant ARUK-PG2017B−10. S.R.C. and I.J.D. were supported by a National Institutes of Health (NIH) research grant R01AG054628, and S.R.C is supported by a Sir Henry Dale Fellowship jointly funded by the Wellcome Trust and the Royal Society (Grant Number 221890/Z/20/Z).

## References

Almstrup, K., Lindhardt Johansen, M., Busch, A.S., Hagen, C.P., Nielsen, J.E., Petersen, J.H., and Juul, A. (2016). Pubertal development in healthy children is mirrored by DNA methylation patterns in peripheral blood. Sci. Rep. 6, 28657.

Baglietto, L., Ponzi, E., Haycock, P., Hodge, A., Bianca Assumma, M., Jung, C.-H., Chung, J., Fasanelli, F., Guida, F., Campanella, G., et al. (2017). DNA methylation changes measured in pre-diagnostic peripheral blood samples are associated with smoking and lung cancer risk. Int. J. Cancer 140, 50–61.

Barker, E.D., Walton, E., and Cecil, C.A.M. (2018). Annual Research Review: DNA methylation as a mediator in the association between risk exposure and child and adolescent psychopathology. J. Child Psychol. Psychiatry 59, 303–322.

Benzeval, M., Davillas, A., Kumari, M., and Lynn, P. (2014). Understanding society: the UK household longitudinal study biomarker user guide and glossary (Institute for Social and Economic Research, University of Essex).

Bose, M., Wu, C., Pankow, J.S., Demerath, E.W., Bressler, J., Fornage, M., Grove, M.L., Mosley, T.H., Hicks, C., North, K., et al. (2014). Evaluation of microarray-based DNA methylation measurement using technical replicates: the Atherosclerosis Risk In Communities (ARIC) Study. BMC Bioinformatics 15, 312.

Boyd, A., Golding, J., Macleod, J., Lawlor, D.A., Fraser, A., Henderson, J., Molloy, L., Ness, A., Ring, S., and Davey Smith, G. (2013). Cohort Profile: The ‘Children of the 90s’—the index offspring of the Avon Longitudinal Study of Parents and Children. Int. J. Epidemiol. 42, 111–127.

Canfield, C.-A., and Bradshaw, P.C. (2019). Amino acids in the regulation of aging and aging-related diseases. Transl. Med. Aging 3, 70–89.

Christiansen, C., Castillo-Fernandez, J.E., Domingo-Relloso, A., Zhao, W., El-Sayed Moustafa, J.S., Tsai, P.-C., Maddock, J., Haack, K., Cole, S.A., Kardia, S.L.R., et al. (2021). Novel DNA methylation signatures of tobacco smoking with trans-ethnic effects. Clin. Epigenetics 13, 36.

Deary, I.J., Gow, A.J., Pattie, A., and Starr, J.M. (2012). Cohort profile: the Lothian Birth Cohorts of 1921 and 1936. Int. J. Epidemiol. 41, 1576–1584.

Dugué, P.-A., English, D.R., MacInnis, R.J., Jung, C.-H., Bassett, J.K., FitzGerald, L.M., Wong, E.M., Joo, J.E., Hopper, J.L., Southey, M.C., et al. (2016). Reliability of DNA methylation measures from dried blood spots and mononuclear cells using the HumanMethylation450k BeadArray. Sci. Rep. 6, 30317.

Elliott, J., and Shepherd, P. (2006). Cohort Profile: 1970 British Birth Cohort (BCS70). Int. J. Epidemiol. 35, 836–843.

Fraser, A., Macdonald-Wallis, C., Tilling, K., Boyd, A., Golding, J., Davey Smith, G., Henderson, J., Macleod, J., Molloy, L., Ness, A., et al. (2013). Cohort Profile: The Avon Longitudinal Study of Parents and Children: ALSPAC mothers cohort. Int. J. Epidemiol. 42, 97–110.

Gaunt, T.R., Shihab, H.A., Hemani, G., Min, J.L., Woodward, G., Lyttleton, O., Zheng, J., Duggirala, A., McArdle, W.L., Ho, K., et al. (2016). Systematic identification of genetic influences on methylation across the human life course. Genome Biol. 17, 61.

Han, L., Zhang, H., Kaushal, A., Rezwan, F.I., Kadalayil, L., Karmaus, W., Henderson, A.J., Relton, C.L., Ring, S., Arshad, S.H., et al. (2019). Changes in DNA methylation from pre-to post-adolescence are associated with pubertal exposures. Clin. Epigenetics 11, 176.

Hannum, G., Guinney, J., Zhao, L., Zhang, L., Hughes, G., Sadda, S., Klotzle, B., Bibikova, M., Fan, J.-B., Gao, Y., et al. (2013). Genome-wide methylation profiles reveal quantitative views of human aging rates. Mol. Cell 49, 359–367.

Hernando-Herraez, I., Evano, B., Stubbs, T., Commere, P.-H., Jan Bonder, M., Clark, S., Andrews, S., Tajbakhsh, S., and Reik, W. (2019). Ageing affects DNA methylation drift and transcriptional cell-to-cell variability in mouse muscle stem cells. Nat. Commun. 10, 4361.

Horvath, S. (2013). DNA methylation age of human tissues and cell types. Genome Biol. 14, R115.

Jones, S., Tillin, T., Park, C., Williams, S., Rapala, A., Al Saikhan, L., Eastwood, S.V., Richards, M., Hughes, A.D., and Chaturvedi, N. (2020). Cohort Profile Update: Southall and Brent Revisited (SABRE) study: a UK population-based comparison of cardiovascular disease and diabetes in people of European, South Asian and African Caribbean heritage. Int. J. Epidemiol. 49, 1441–1442e.

Maddock, J., Castillo-Fernandez, J., Wong, A., Cooper, R., Richards, M., Ong, K.K., Ploubidis, G.B., Goodman, A., Kuh, D., Bell, J.T., et al. (2019). DNA methylation age and physical and cognitive ageing. J. Gerontol. Ser. A.

McCartney, D.L., Zhang, F., Hillary, R.F., Zhang, Q., Stevenson, A.J., Walker, R.M., Bermingham, M.L., Boutin, T., Morris, S.W., Campbell, A., et al. (2019). An epigenome-wide association study of sex-specific chronological ageing. Genome Med. 12, 1.

Mendelsohn, A.R., and Larrick, J.W. (2017). Epigenetic Drift Is a Determinant of Mammalian Lifespan. Rejuvenation Res. 20, 430–436.

Milnik, A., Vogler, C., Demougin, P., Egli, T., Freytag, V., Hartmann, F., Heck, A., Peter, F., Spalek, K., Stetak, A., et al. (2016). Common epigenetic variation in a European population of mentally healthy young adults. J. Psychiatr. Res. 83, 260–268.

Min, J.L., Hemani, G., Davey Smith, G., Relton, C., and Suderman, M. (2018). Meffil: efficient normalization and analysis of very large DNA methylation datasets. Bioinforma. Oxf. Engl. 34, 3983–3989.

Mulder, R.H., Neumann, A., Cecil, C.A.M., Walton, E., Houtepen, L.C., Simpkin, A.J., Rijlaarsdam, J., Heijmans, B.T., Gaunt, T.R., Felix, J.F., et al. (2021). Epigenome-wide change and variation in DNA methylation in childhood: trajectories from birth to late adolescence. Hum. Mol. Genet. 30, 119–134.

Oh, E.S., and Petronis, A. (2021). Origins of human disease: the chrono-epigenetic perspective. Nat. Rev. Genet. 22, 533–546.

Platt, L., Knies, G., Luthra, R., Nandi, A., and Benzeval, M. (2020). Understanding Society at 10 years. Eur. Sociol. Rev. 36, 976–988.

Power, C., and Elliott, J. (2006). Cohort profile: 1958 British birth cohort (National Child Development Study). Int. J. Epidemiol. 35, 34–41.

Relton, C.L., Gaunt, T., McArdle, W., Ho, K., Duggirala, A., Shihab, H., Woodward, G., Lyttleton, O., Evans, D.M., Reik, W., et al. (2015). Data Resource Profile: Accessible Resource for Integrated Epigenomic Studies (ARIES). Int. J. Epidemiol. 44, 1181–1190.

Rubio-Aparicio, M., López-López, J.A., Viechtbauer, W., Marín-Martínez, F., Botella, J., and Sánchez-Meca, J. (2020). Testing Categorical Moderators in Mixed-Effects Meta-analysis in the Presence of Heteroscedasticity. J. Exp. Educ. 88, 288–310.

Ryan, J., Wrigglesworth, J., Loong, J., Fransquet, P.D., and Woods, R.L. (2020). A Systematic Review and Meta-analysis of Environmental, Lifestyle, and Health Factors Associated With DNA Methylation Age. J. Gerontol. Ser. A 75, 481–494.

Saxton, R.A., and Sabatini, D.M. (2017). mTOR signaling in growth, metabolism, and disease. Cell 168, 960–976.

Slieker, R.C., van Iterson, M., Luijk, R., Beekman, M., Zhernakova, D.V., Moed, M.H., Mei, H., Van Galen, M., Deelen, P., and Bonder, M.J. (2016). Age-related accrual of methylomic variability is linked to fundamental ageing mechanisms. Genome Biol. 17, 1–13.

Smith, Z.D., and Meissner, A. (2013). DNA methylation: roles in mammalian development. Nat. Rev. Genet. 14, 204–220.

Snir, S., Farrell, C., and Pellegrini, M. (2019). Human epigenetic ageing is logarithmic with time across the entire lifespan. Epigenetics 14, 912–926.

Staley, J.R., Windmeijer, F., Suderman, M., Lyon, M.S., Smith, G.D., and Tilling, K. (2020). A robust mean and variance test with application to high-dimensional phenotypes. BioRxiv 2020.02.06.926584.

Sugden, K., Hannon, E.J., Arseneault, L., Belsky, D.W., Corcoran, D.L., Fisher, H.L., Houts, R.M., Kandaswamy, R., Moffitt, T.E., Poulton, R., et al. (2020). Patterns of Reliability: Assessing the Reproducibility and Integrity of DNA Methylation Measurement. Patterns N. Y. N 1, 100014.

Talens, R.P., Christensen, K., Putter, H., Willemsen, G., Christiansen, L., Kremer, D., Suchiman, H.E.D., Slagboom, P.E., Boomsma, D.I., and Heijmans, B.T. (2012). Epigenetic variation during the adult lifespan: cross-sectional and longitudinal data on monozygotic twin pairs. Aging Cell 11, 694–703.

Taylor, A.M., Pattie, A., and Deary, I.J. (2018). Cohort Profile Update: The Lothian Birth Cohorts of 1921 and 1936. Int. J. Epidemiol. 47, 1042–1042r.

Tillin, T., Forouhi, N.G., McKeigue, P.M., Chaturvedi, N., and SABRE Study Group (2012). Southall And Brent REvisited: Cohort profile of SABRE, a UK population-based comparison of cardiovascular disease and diabetes in people of European, Indian Asian and African Caribbean origins. Int. J. Epidemiol. 41, 33–42.

Valiathan, R., Ashman, M., and Asthana, D. (2016). Effects of Ageing on the Immune System: Infants to Elderly. Scand. J. Immunol. 83, 255–266.

Verdi, S., Abbasian, G., Bowyer, R.C.E., Lachance, G., Yarand, D., Christofidou, P., Mangino, M., Menni, C., Bell, J.T., Falchi, M., et al. (2019). TwinsUK: The UK Adult Twin Registry Update. Twin Res. Hum. Genet. Off. J. Int. Soc. Twin Stud. 22, 523–529.

Wang, D., Liu, X., Zhou, Y., Xie, H., Hong, X., Tsai, H.-J., Wang, G., Liu, R., and Wang, X. (2012). Individual variation and longitudinal pattern of genome-wide DNA methylation from birth to the first two years of life. Epigenetics 7, 594–605.

Yashin, A.I., Wu, D., Arbeev, K.G., and Ukraintseva, S.V. (2010). Joint influence of small-effect genetic variants on human longevity. Aging 2, 612–620.

Yuan, T., Jiao, Y., de Jong, S., Ophoff, R.A., Beck, S., and Teschendorff, A.E. (2015). An integrative multi-scale analysis of the dynamic DNA methylation landscape in aging. PLoS Genet 11, e1004996.

Yusipov, I., Bacalini, M.G., Kalyakulina, A., Krivonosov, M., Pirazzini, C., Gensous, N., Ravaioli, F., Milazzo, M., Giuliani, C., Vedunova, M., et al. (2020). Age-related DNA methylation changes are sex-specific: a comprehensive assessment. Aging 12, 24057–24080.

