## Supplemental Materials for "Characterizing the human methylome across the life course: findings from eight UK-based studies"

**SM 1 Methods**

*SM 1.1 Sample description*

ALSPAC

Pregnant women resident in Avon, UK with expected dates of delivery 1st April 1991 to 31st December 1992 were invited to take part in the study. The initial number of pregnancies enrolled is 14,541 (for these at least one questionnaire has been returned or a “Children in Focus” clinic had been attended by 19/07/99). Of these initial pregnancies, there was a total of 14,676 foetuses, resulting in 14,062 live births and 13,988 children who were alive at 1 year of age. For more information, see references (Boyd et al., 2013; Fraser et al., 2013; Northstone et al., 2019). A sub-set of 1018 mother-offspring pairs were included in Accessible resource for integrated epigenomics studies (ARIES) and were selected based on availability of DNA samples at two time-points for the mother (at an antenatal clinic and at a follow-up clinic when their offspring were mean age 15.5 years) and three time points for the offspring [at birth, childhood (mean age 7.5 years) and adolescence (mean age 15.5 years)]. Cord and peripheral blood samples (whole blood or buffy coat) were collected according to standard procedures. Some of the cord blood samples were obtained from blood spots. Following extraction, DNA was bisulfite-converted using the Zymo EZ DNA MethylationTM kit (Zymo, Irvine, CA). The arrays were scanned using an Illumina iScan and initial quality review was assessed using GenomeStudio. For more information, see reference (Relton et al., 2015).

In addition to these data, we also included DNA methylation data for the fathers (mean age 53 years) and data at five additional time points for the offspring [4, 5, 7, 10 and 24 years]. Please note that the study website contains details of all the data that is available through a fully searchable data dictionary and variable search tool (<http://www.bristol.ac.uk/alspac/researchers/our-data/>). Study data were collected and managed using REDCap electronic data capture tools (Harris et al., 2009).

Ethical approval for the study was obtained from the ALSPAC Ethics and Law Committee and the Local Research Ethics Committees. Informed consent for the use of data collected via questionnaires and clinics was obtained from participants following the recommendations of the ALSPAC Ethics and Law Committee at the time. Consent for biological samples has been collected in accordance with the Human Tissue Act (2004).

1958 National Child Development Study

The 1958 British birth cohort or National Child Development Study (NCDS) (Power & Elliott, 2006) comprises >17,000 people born in England, Scotland and Wales in a single week of 1958.

Blood samples from 240 participants were processed to extract DNA and characterise DNA methylation profiles using the Illumina Infinium MethylationEPIC BeadChips, as previously described (Christiansen et al., 2021; Maddock et al., 2019).

1970 British Cohort Study

The 1970 British Birth Cohort Study (BCS70) (Elliott & Shepherd, 2006) is an ongoing, multidisciplinary, longitudinal study. It takes as its subjects all those currently living in England, Scotland, and Wales who were born in a single week of 1970.

For this study, 235 blood samples were extracted when participants were 46 years of age and processed using the Illumina EPIC DNA methylation array (Christiansen et al., 2021).

TwinsUK

TwinsUK is a nationwide registry of adult twins that started in 1992. The registry has recruited more 14,000 volunteers whose majority of participants are adult females of European descent (Verdi et al., 2019).

For this study we included 120 female MZ twins with existing whole blood DNA methylation profiles without selecting for a particular disease (Christiansen et al., 2021; Maddock et al., 2019). DNA for methylation assessment was extracted from whole blood and stored in EDTA tubes. The Illumina Infinium MethylationEPIC BeadChip (Illumina Inc, CA, USA) was used to measure DNA methylation levels.

Ethical approval was granted by the National Research Ethics Service London-Westminster, the St Thomas’ Hospital Research Ethics Committee (EC04/015 and 07/H0802/84). All research participants have signed informed consent prior to taking part in any research activities.

Understanding Society

The goal of *Understanding Society* is to assess long-term and short-term effects of social and economic change on a variety of outcomes. Social and economic data of 1193 individuals were recorded through questionnaires and additional information including biomarker data and genotyping micro-arrays have also been obtained. Biomarker and relevant questionnaire data are available at <https://www.understandingsociety.ac.uk/about/health/data> upon request. 500Ng of whole blood DNA from each individual was treated with sodium bisulfite using the EZ96 DNA methylation kit (Zymo Research, CA, USA) following manufacturer’s standard protocol. DNA methylation intensities were assess using Illumina Infinium HumanMethylationEPIC BeadChips (Illumina Inc, CA, USA) in the Laboratory of Professor Jon Mill (University of Exeter). DNA methylation levels were assessed on an Illumina HiScan System (Illumina). For more information, see reference (Gorrie-Stone et al., 2019).

DNA methylation data was pre-processed, as further described in reference (Mansell et al., 2019). Briefly, raw signal intensity data were processed from idat files through a standard pipeline using the bigmelon and wateRmelon packages in R. A number of quality control steps were performed to these data prior to normalization. First, outlier samples were identified using principal component analysis and mahalanobis distance equivalents, second, successful bisulphite conversion was confirmed using control probes, third the ages of the samples were estimated using the Horvath Epigenetic Clock algorithm and compared to reported age at sampling, and fourth visualisation of principal components. These data were then normalized using the dasen method, which performs background adjustment and between-sample quantile normalization of methylated (M) and unmethylated (U) intensities separately for Type I and Type II probes. A second round of sample filtering was then performed excluding samples that were either dramatically altered as a result of normalisation or samples that had > 1% of sites with detection p-value > 0.05. DNAm sites were filtered to exclude those with a bead count < 3 or > 1% of samples with detection p-value > 0.05. The raw DNAm data of the final sample set was then re-normalized with the dasen method.

SABRE

Samples were derived from the extensively characterised population-based Southall And Brent Revisited (SABRE) cohort (Tillin et al., 2012). The SABRE cohort includes 1,711 first generation South Asian migrants and 1,762 European origin individuals aged between 40-69 living in west London, UK, with Initial investigations carried out between 1988 and 1991 and follow up at almost 20 years later. DNA was extracted from peripheral blood samples collected at baseline and follow up visits (Jones et al., 2020). Samples selected for methylation analysis were those who were male, self-reported their ethnic group as “Indian Asian” or “European” and who had a good quality DNA sample available. Ethical approval was granted by Fulham Research Ethics Committee (14/LO/0108) and all participants provided written informed consent.

Genomic DNA (500 ng) was bisulphite modified using an EZ DNA methylation kit (Zymo Research, Orange, CA, USA). The protocol was as described by the manufacturer, utilising the alternative incubation conditions recommended when using Illumina Infinium Methylation Arrays. Genome-wide methylation was measured using the Illumina HumanMethylation450 BeadChip (Illumina, San Diego, CA, USA) following the manufacturer’s protocol with no modifications. The arrays were scanned using an Illumina iScan with software version 3.3.28. Initial quality control of sample data was conducted using GenomeStudio version 2011.1 (Illumina, San Diego, CA, USA) to determine the status of staining, extension, hybridisation, target removal, bisulphite conversion, specificity, non-polymorphic and negative controls.

Lothian Birth Cohort 1921

Data were from the Lothian Birth Cohort 1921 (LBC1921), which is the basis of a longitudinal study of aging (Deary et al., 2012; Taylor et al., 2018). Participants were born in 1921 and most completed a cognitive ability test at about the age of 11 years in the Scottish Mental Survey 1932 (SMS1932. The LBC1921 study attempted to follow up individuals who might have completed the SMS1932 and resided at about the age of 79 years in the Lothian region (Edinburgh and its surrounding areas) of Scotland; 550 people (n=234, 43% men) were successfully traced and participated in the study from the age of 79 years. To date, there have been four additional follow-up waves at average ages of 83, 87, 90, and 92 years. DNA methylation used for analyses in the current study was measured in three of these five waves, with subjects at an average age of 79, 87 and 90. DNA methylation was measured using the Illumina HumanMethylation450BeadChips. Ethics permission for the LBC1921 was obtained from the Lothian Research Ethics Committee (Wave 1: LREC/1998/4/183). Written informed consent was obtained from all subjects. For further information, see (Shah et al., 2014; Zhang et al., 2018).

Lothian Birth Cohort 1936

The Lothian Birth Cohort 1936 (LBC1936) was established to study cognitive aging in surviving members of the 1947 Scottish Mental Survey. All participants were born in 1936. 1091 community-dwelling people were recruited aged around 70 years, mostly from the Edinburgh area of Scotland. Initially, whole blood was obtained in 1004 participants from samples collected at mean age 70 years (wave 1). There have been four additional follow-up waves at average ages of 73, 76, 79, and 82 years. Data from the first three waves (ages 70-76) were used in the current study. DNA methylation was measured using the Illumina HumanMethylation450BeadChips. Ethical approval was obtained from the Multi-Centre Ethics Committee for Scotland and Lothian Research Ethics Committee. All subjects provided written informed consent. For further information, see (Shah et al., 2014; Zhang et al., 2018).

*SM 1.2 Cross-cohort harmonization and normalization*

SM Figure 1 shows a schematic work flow used to harmonize and normalize DNA methylation samples across all cohorts (shown as “EPIC_1”, “EPIC_2”, etc). We followed three steps. In step 1, qc.object files were derived for each cohort and samples failing quality control were excluded at this step (e.g. due to genotype or gender mismatches, low detection scores, low number of beads, methylated/unmethylated ratio). Since these qc.object files capture only technical variation and data summaries, the information they contain cannot be used to identify individuals, thus removing any objections to them being shared centrally. In step 2, all qc.object files were pooled, normalized and returned to each cohort. In step 3, raw probe intensities for each sample were then adjusted to conform to its normalized data summary. At this step, CpG sites failing quality control were excluded (e.g. low detection scores, low number of beads).

Of note, functional normalization preserves mean and variance structure of the data. Briefly, functional normalization begins by calculating percentiles of the raw DNA methylation levels for each sample. The values for each percentile are then adjusted across samples to remove control probe variation. Finally, the raw DNA methylation levels are adjusted for each sample so that their percentiles match its adjusted percentiles.

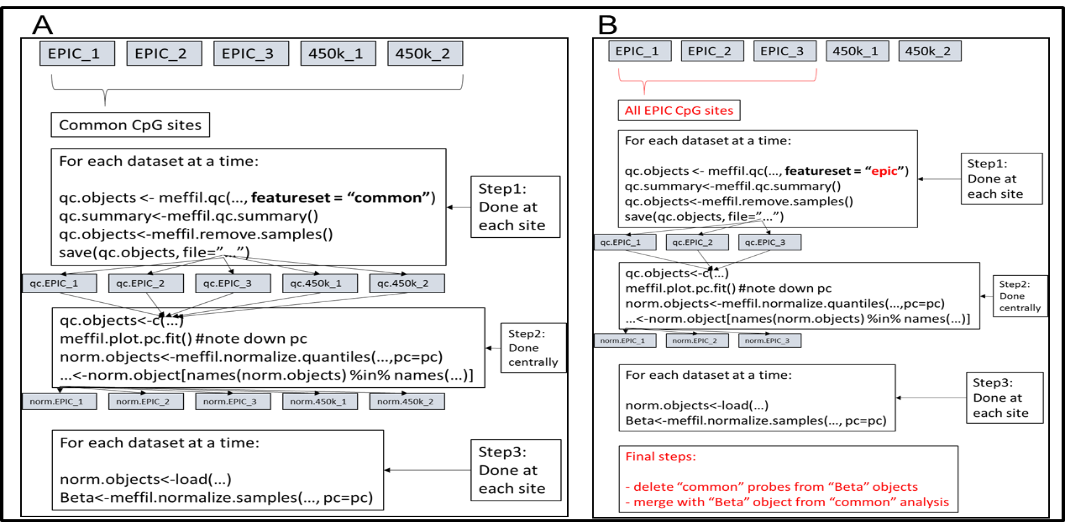
As we included data from both EPIC and 450k arrays, this workflow was repeated for CpG sites i) common to both arrays (SM Figure 1A), ii) all CpG sites on the EPIC array (SM Figure 1B) and iii) all CpG sites on the 450k array (not shown; as SM Figure 1B but “featureset = ‘450k’ ”). For each cohort, a final dataset was created by taking the “common probes” set as a base and adding to this base, probes unique to the EPIC and to the 450k array, respectively. This way, a given probe was normalized against the largest possible set of data (e.g. a probe present on both the EPIC and 450k array was normalized against all datasets while a probe unique to the EPIC array was normalized against all EPIC datasets, only). Detailed scripts are available at <https://github.com/stegosaurusrox/Lifecourse_Methylome>.

*SM Figure 1*. Normalization and harmonization workflow for CpG sites A) common to both the EPIC and 450k Illumina array and B) for CpG sites on the EPIC array. Step B was repeated for 450k sites by setting featureset to “450k” (not shown).

*SM 1.3 Analysis Flow diagram*
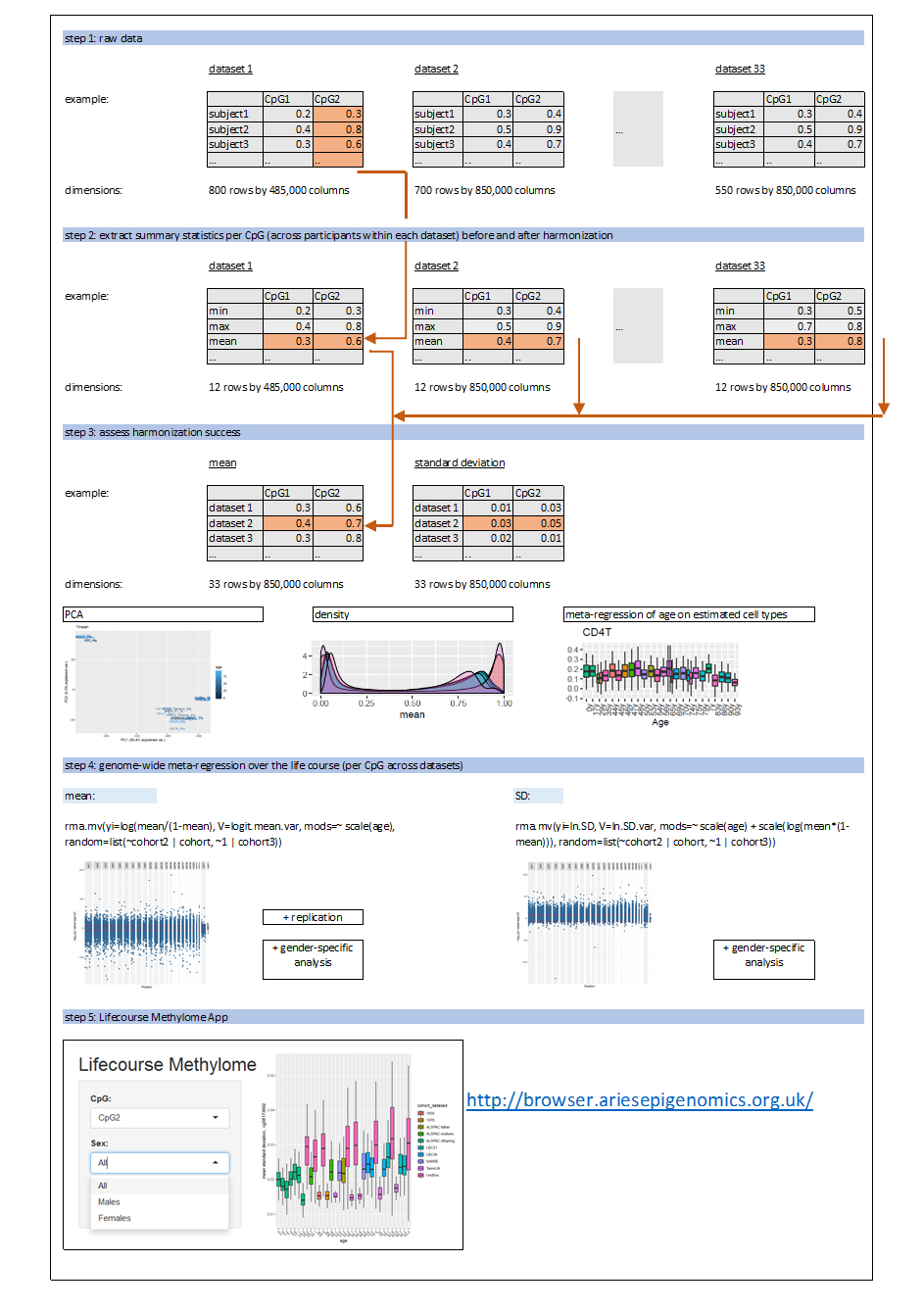

*SM Figure 2*. Analysis flow diagram.

*SM 1.4 Assessment of harmonization success*

We assessed the importance of cross-cohort normalization by comparing DNA methylation summary data before and after harmonization i) using principal component analysis, ii) inspecting density curves across statistics and datasets and iii) by assessing the change in study heterogeneity observed in a meta-regression of age on methylation-based cell type composition.

Using principal component analysis across mean methylation and its standard deviation, we observed two distinct clusters of cohorts before harmonization, especially for mean methylation (SM Figure 3). We suspect these clusters reflect different preprocessing procedures, as the observed clustering was substantially reduced after data harmonization. *Understanding Society* data, which was not harmonized with the other datasets, did not cluster separately for SD methylation and was only slightly apart from the other cohorts for mean methylation.

Furthermore, we inspected density curves across all 33 datasets. For comparative purposes, our main analysis focussed on summary statistics based on CpGs *shared* across all dataset (CpG n=453,008). As before, cross-cohort differences were clearly visible before harmonization, especially for mean methylation, and reduced after harmonization (SM Figure 4).

Last, we assessed the amount of residual heterogeneity after modelling the effect of age on the six different cell types (Bcells, CD4T, CD8T, Granulocytes, monocytes and natural killer cells, but not nucleated red blood cells, as these were only available at birth) across the life course. On average, heterogeneity decreased after harmonization (mean I^2^_pre_ = 28.82 versus mean I^2^_post_ = 14.95; SM Table 2). Taken together, these observations indicate substantial differences in raw DNA methylation data patterns between cohorts, which we were able to minimize through our harmonization protocol.

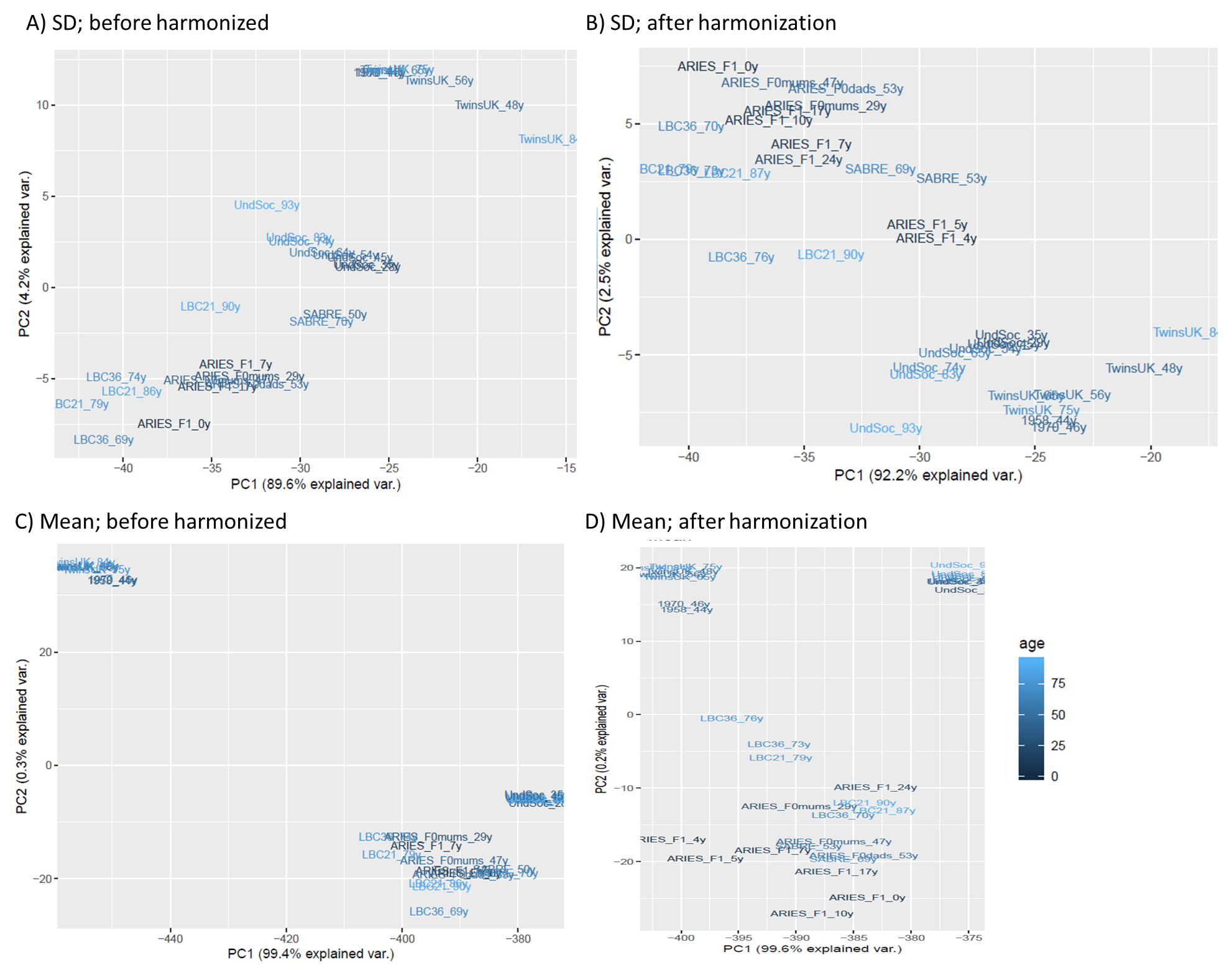

*SM Figure 3*. Cohort clustering of A-B) SD methylation and C-D) mean methylation, visible before harmonization (A/C) and after harmonization (B/D).

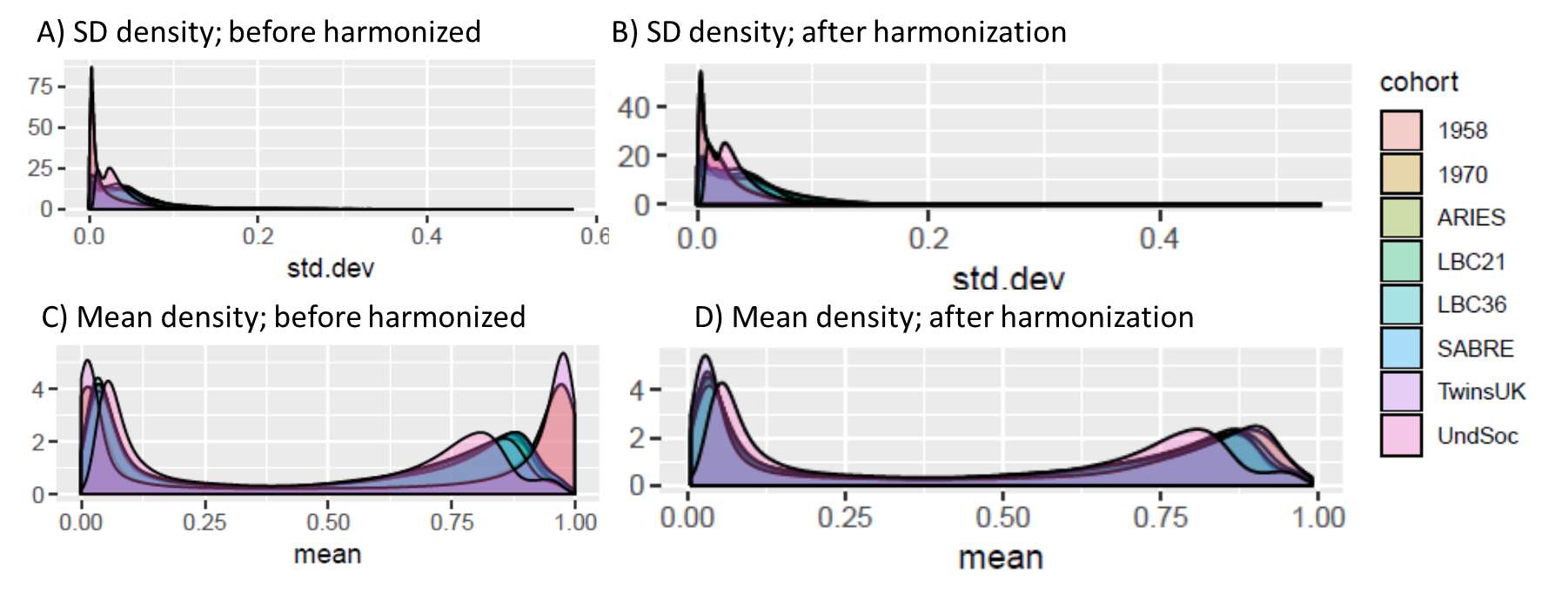

*SM Figure 4*. Density plots of SD and SD methylation before and after harmonization.

*SM 1.5 Change of SD and mean DNA methylation over the life course*

To assess common linear trends in SD and mean DNA methylation per CpG over the life course across cohorts, we carried out meta-regressions of DNA methylation on age using summary data. We allowed for the correlation between related (e.g. familiar) or repeated time points in the same cohort. We also allowed for independent residual variation for each measurement. Analyses were carried out using the metafor R package (Viechtbauer, 2010).

To identify age-related trends of SD DNA methylation, we applied the following regression model:

$$log\left( S_{ijk} \right)= \beta_{0}+\beta_{1}age_{ijk}+{\beta_{2}log\left( M_{ijk}(1-M_{ijk}) \right)+u}_{i}+u_{ij}+\varepsilon_{ijk}+e_{ijk},u_{i}\sim N\left( 0,\rho\tau^{2} \right),u_{ij}\sim N\left( 0,\left( 1-\rho\right)\tau^{2} \right),\varepsilon_{ijk}\sim N\left( 0,\sigma^{2} \right),e_{ijk}\sim N\left( 0,\frac{1}{{2(n}_{ijk}-1)} \right).$$

where $M_{ijk}$ and $S_{ijk}$ are the sample mean and standard deviation, respectively, of the DNA methylation at a particular CpG site at time point $k$ for cohort $j$ in study $i$. We applied a log transformation to the standard deviation, so that it is not restricted to be positive. The sample size, $n_{ijk}$, and average age, $age_{ijk}$, are dependent on the CpG site. Random effects are present at three levels in the model: $u_{i}$ represents a study effect, $u_{ij}$ represents a cohort effect, and $\varepsilon_{ijk}$ represents the usual residual at each time point. These random effects have a covariance structure parametrised by $\tau^{2}$, the variance of the cohort effects, $\rho$, the correlation between cohort effects for cohorts in the same study, and $\sigma^{2}$, the residual variance. The measurement error, $e_{ijk}$, is likely to be larger in smaller cohorts and also when the cohort sample standard deviation is larger. Therefore, the measurement error has a variance that depends on the cohort sample size and sample standard deviation.

In this model, we controlled for effects of mean methylation due to the assumption that the variance is influenced by the mean (but not vice versa). This is represented by the term $log\left( M_{ijk}(1-M_{ijk}) \right)$, chosen as the standard deviation of levels representing a proportion is often expected to have a parabolic relationship with the mean.^10^ As expected, the linear trend between the mean and standard deviation increased substantially after log transformation (mean R^2^_raw_ across 33 datasets = 6% versus R^2^_transf_ = 78%; SM Figure 5A and 5B as well as SM Table 3).

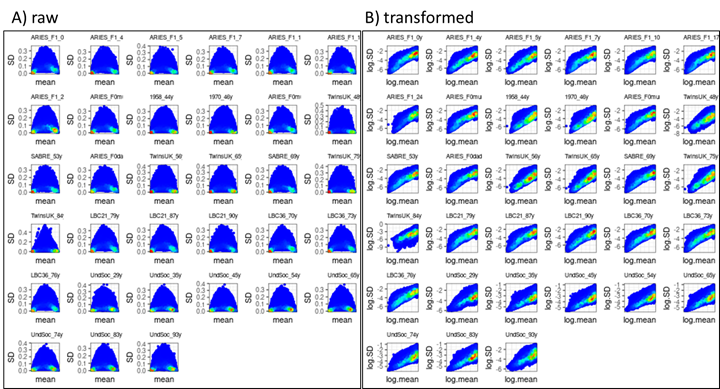

*SM Figure 5*. Density plots per dataset of A) untransformed means versus SD and B) log-transformed means [$log\left( M_{ijk}(1-M_{ijk}) \right)$] versus log(SD) across all CpGs.

To identify age-related trends in *mean* DNA methylation, we used the following model:

$$log\left( \frac{M_{ijk}}{1-M_{ijk}} \right)= \beta_{0}+\beta_{1}age_{ijk}+u_{i}+u_{ij}+\varepsilon_{ijk}+e_{ijk},u_{i}\sim N\left( 0,\rho\tau^{2} \right),u_{ij}\sim N\left( 0,\left( 1-\rho\right)\tau^{2} \right),\varepsilon_{ijk}\sim N\left( 0,\sigma^{2} \right),e_{ijk}\sim N\left( 0,\frac{S_{ijk}^{2}}{n_{ijk}\left( M_{ijk} \right)^{2}\left( 1-M_{ijk} \right)^{2}} \right),$$

This model uses the same notation as the model for SD DNA methylation (see above). We applied a logit transformation on mean methylation levels, because these levels represent a proportion of total methylation.

In sensitivity analyses, we additionally controlled for cell type composition using the cell type reference panel developed by Reinius et al. (2012) and the algorithms developed by Houseman (2012). Of note, models for n=116,771 and n=75,495 CpGs did not converge in these cell-type adjusted analyses.

*SM 2.1 QQ plots and correction for cellular composition*
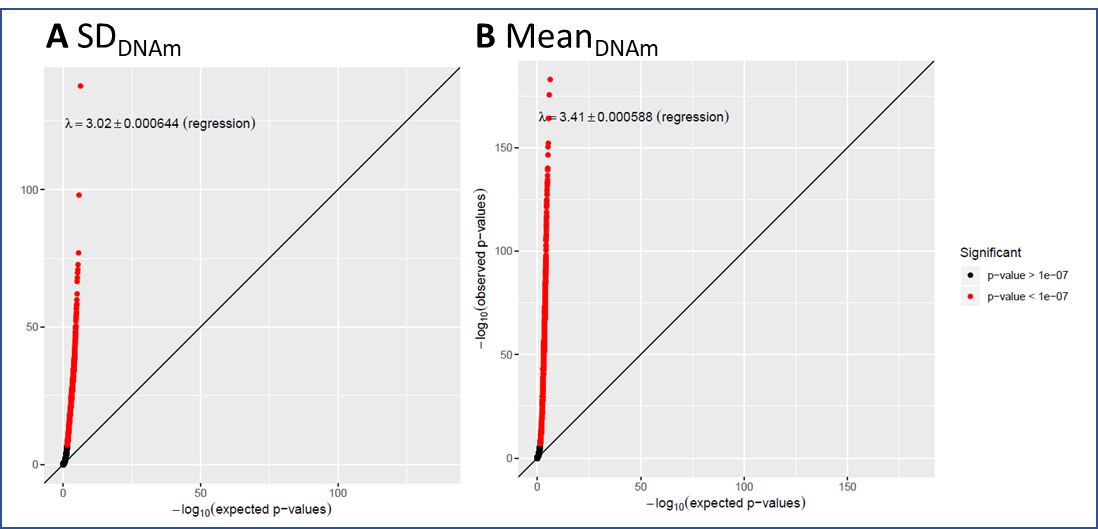

*SM Figure 6*. QQ plots of age effects on A) SD_DNAm_ and B) mean_DNAm_.

SD_DNAm_: We carried forward 29,212 CpGs, which associated with age, in a follow-up analysis additionally controlling for cellular composition. As before, the majority of the associations (99.61%) were positive, indicating an increase in CpG-specific variability over the life course even after cell type correction. We observed a slightly attenuated fold increase of 1.23 in SD_DNAm_ per year over and above any increase expected from concurrent changes in mean methylation or cellular composition. Residual heterogeneity remained in only 26.27% of CpG sites (mean tau^2^=0.008).

Mean_DNAm_: For these analyses, we carried forward 33,730 CpGs, which associated with age. Correcting for cellular composition, a larger proportion of associations than originally reported (81.02% versus 64.33%) were negatively associated with age, indicating a widespread decrease in CpG-specific mean levels over the life course over and above concurrent changes in cellular composition. However, we observed an attenuation of effect estimates from originally 3% in infancy to 0.14%. Residual heterogeneity remained in only 31.34% of CpG sites (mean tau^2^=0.005).

*SM 2.2 Replication*

The proportion of age-associated mean_DNAm_ CpGs, we identified, was much smaller than reported elsewhere (e.g. 3.75% versus 18.96% in McCartney et al. (2019)). We therefore assessed how well our findings of age effects on mean_DNAm_ levels replicated, using EWAS summary data from GenerationScotland (McCartney et al., 2019), which were based on an unrelated cohort of 2,586 individuals between the ages of 18 and 87 years (here, referred to as replication.1) and a further cohort of 4,450 individuals between the ages of 18 and 93 years (here, referred to as replication.2).

Of the n=33,730 CpG sites, which we identified as associated with age (here, referred to as discovery sample), n=29,787 were present in both replication datasets. Effects of our meta-regression and those reported in the replication datasets correlated very highly (replication.1: rho=0.93; replication.2: rho=0.92; SM Figure 7A-B). Eighty-nine percent of the probes with significant mean_DNAm_ age effects in the discovery data also passed a genome-wide threshold in the replication.1 data, while 96% of probes did so in the replication.2 data, supporting our initial findings. Furthermore, the proportion of probes with decreasing mean_DNAm_ over time was largely consistent across studies (64.33% versus 55.66% in McCartney et al. (2019)).

We repeated these analyses in the n=33,730 CpG sites, which we identified as associated with age, additionally controlling for cellular composition. Correlations between effect estimates of our meta-regression (which now included cell-type correlation) and those reported in the replication datasets were attenuated, but still strong (replication.1: rho=0.74; replication.2: rho=0.81; SM Figure 7C-D). The reverse was also true: regression coefficients of age-associated CpG sites reported in McCartney et al. correlated strongly with our effect estimates (replication.1: rho=0.77; replication.2: rho=0.83).

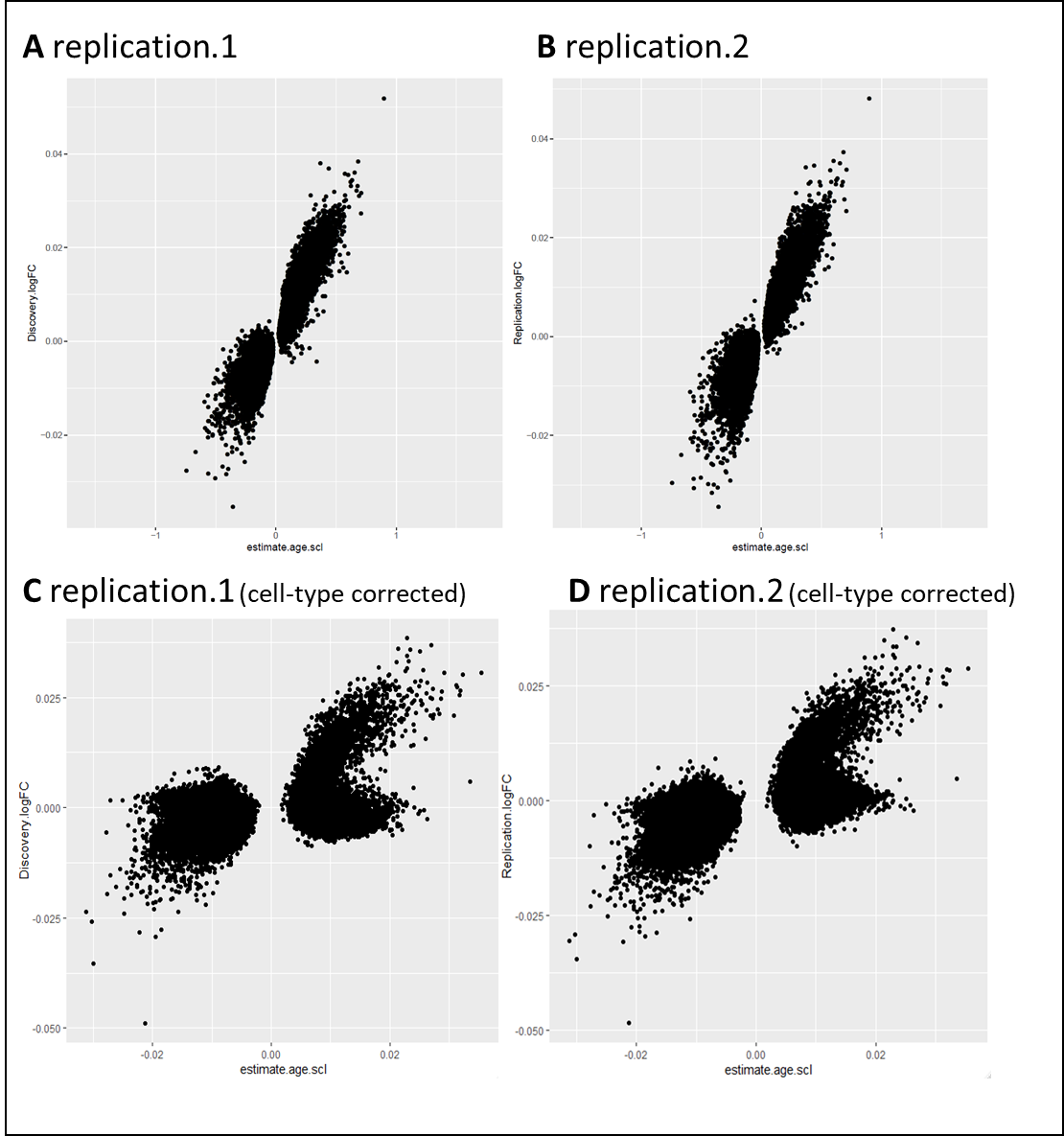

*SM Figure 7*. Replication of age effects on mean_DNAm_ (upper panel: uncorrected effect estimates; lower panel: cell-type corrected effect estimates), using data from A and C) a cohort of 2,586 individuals between the ages of 18 and 87 years (replication.1) and B and D) a cohort of 4,450 individuals between the ages of 18 and 93 years (replication.2).

*SM 2.3 Association between absolute levels of SD methylation or its mean (SD_DNAm_ and mean_DNAm_) and the age effect on these levels (age~SD_DNAm_ and age~mean_DNAm_)*

Correlating absolute levels of SD (the intercept - or β_0_ - in equation 1 below) and the age effect on SD (β_1_) indicated a moderate positive association (SM Figure 8A). For mean methylation (equation 2), this association was slightly negative (SM Figure 8B).

1) $log\left( S_{ijk} \right)= \beta_{0}+\beta_{1}age_{ijk}+{\beta_{2}log\left( M_{ijk}\left( 1-M_{ijk} \right) \right)+u}_{i}+u_{ij}+\varepsilon_{ijk}+e_{ijk},$
2) $log\left( \frac{M_{ijk}}{1-M_{ijk}} \right)= \beta_{0}+\beta_{1}age_{ijk}+u_{i}+u_{ij}+\varepsilon_{ijk}+e_{ijk},$

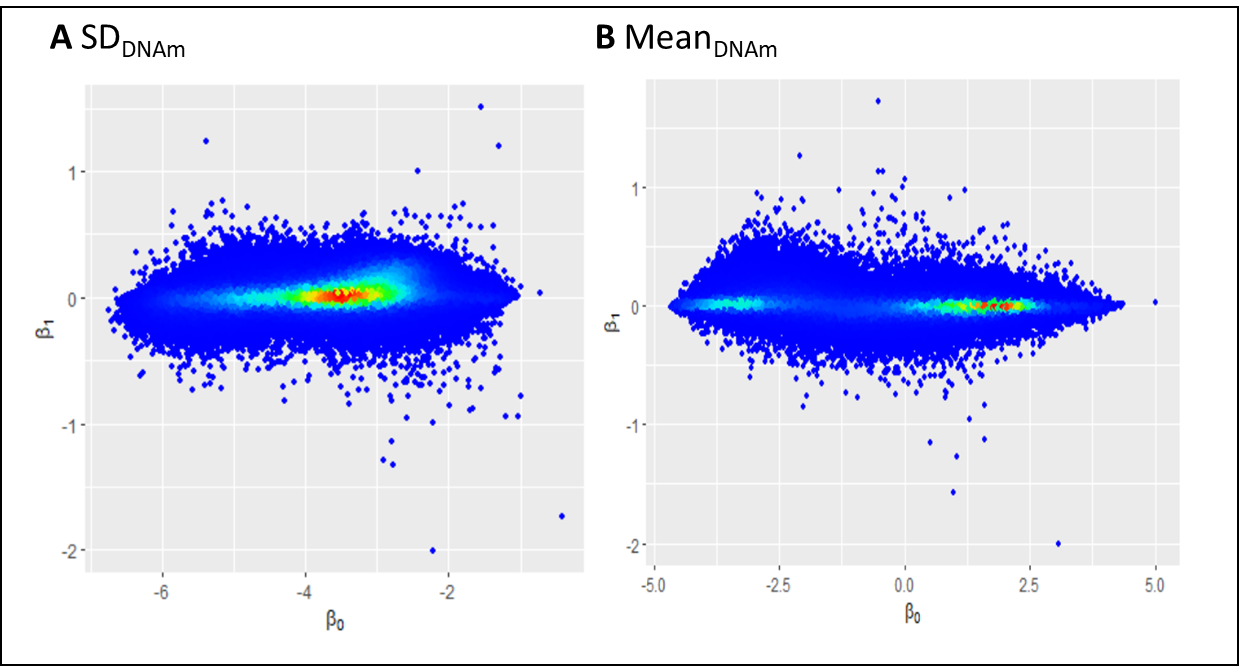

*SM Figure 8*. Density plots for the association between absolute levels of A) SD or B) mean methylation (β_0_; x-axis) and the age effect on these levels (β_1_; y-axis).

*SM 2.4 Associations between age effects and reliability*

To assess the degree to which our findings were influenced by low probe-specific reliability, we correlated reliability measures, as reported in Sugden et al. (2020), with the age effects on mean_DNAm_ and SD_DNAm_. Within each cohort, we could replicate previous findings (SM Figure 9A and 9B), but our meta-analytical results were largely independent of these effects (SM Figure 9C and 9D).

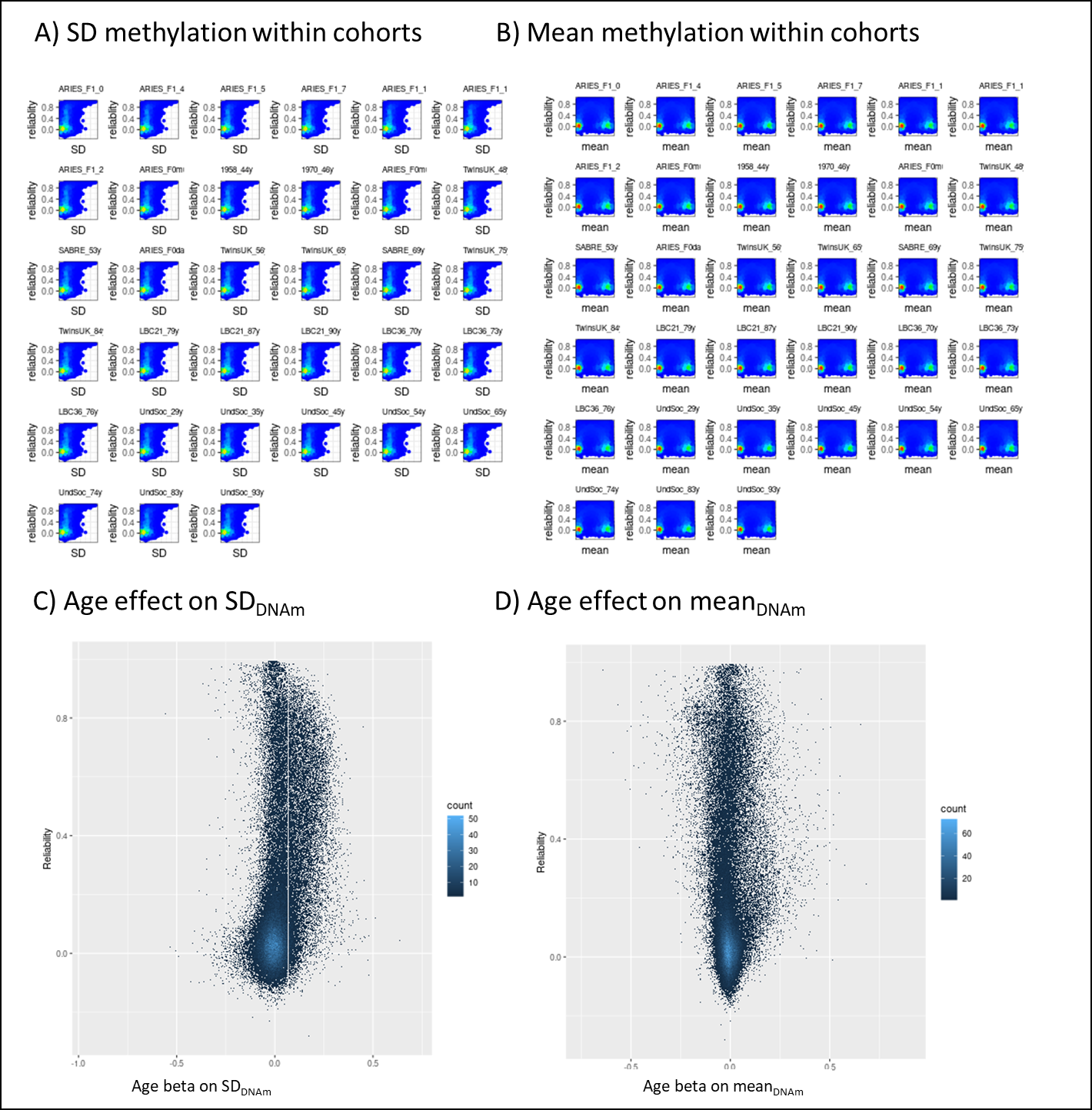

*SM Figure 9*. Correlations between inter-array probe-specific reliability and A) within-cohort SD methylation and B) within-cohort mean methylation, as well as meta-regression age effect on C) cross-cohort SD_DNAm_ and D) cross-cohort mean_DNAm_.

*SM 2.5 Sex-specific effects*

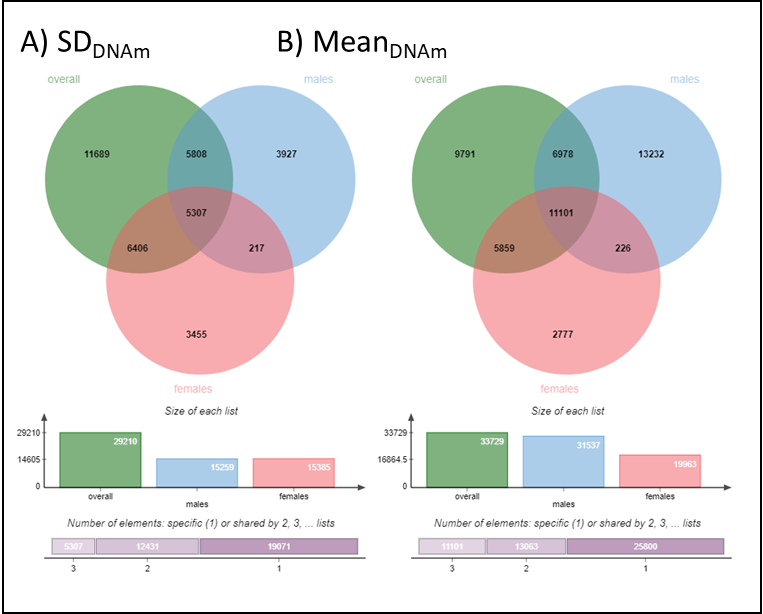

*SM Figure 10*. Number of unique and overlapping CpG sites at genome-wide significance (10^-7^) overall and in males and females for A) SD_DNAm_ and B) mean_DNAm_.

*SM 2.6 EWAS catalogue*

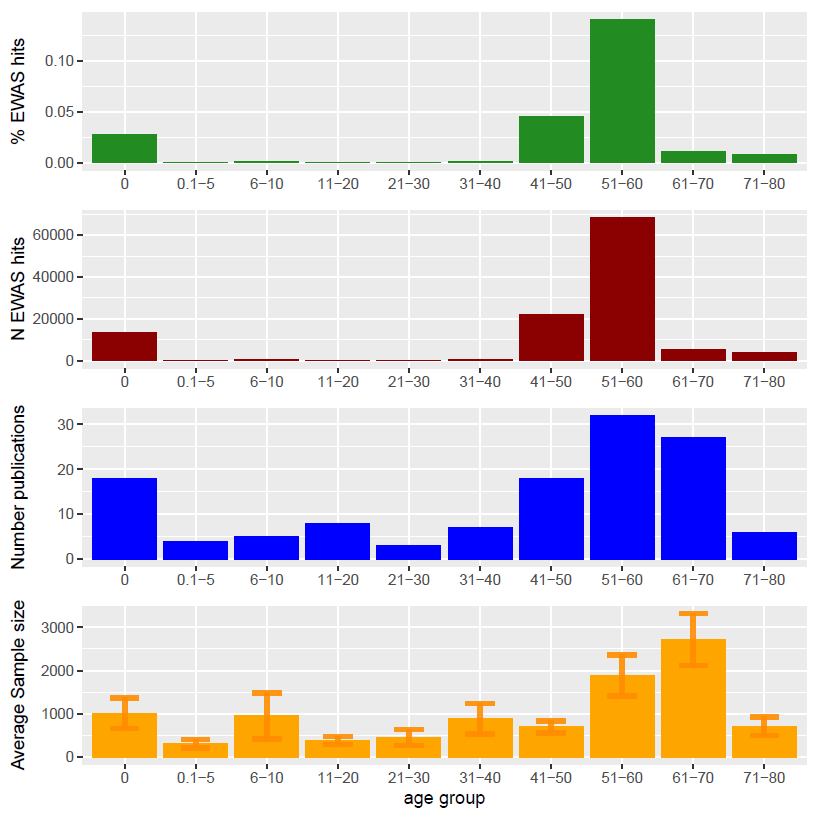

*SM Figure 11*. Overview of content listed on the EWAS catalogue, reduced to epigenome-wide CpG sites reported in blood tissue.

*SM 2.7 Developmental enrichment of trait-associated CpGs: mean_DNAm_*

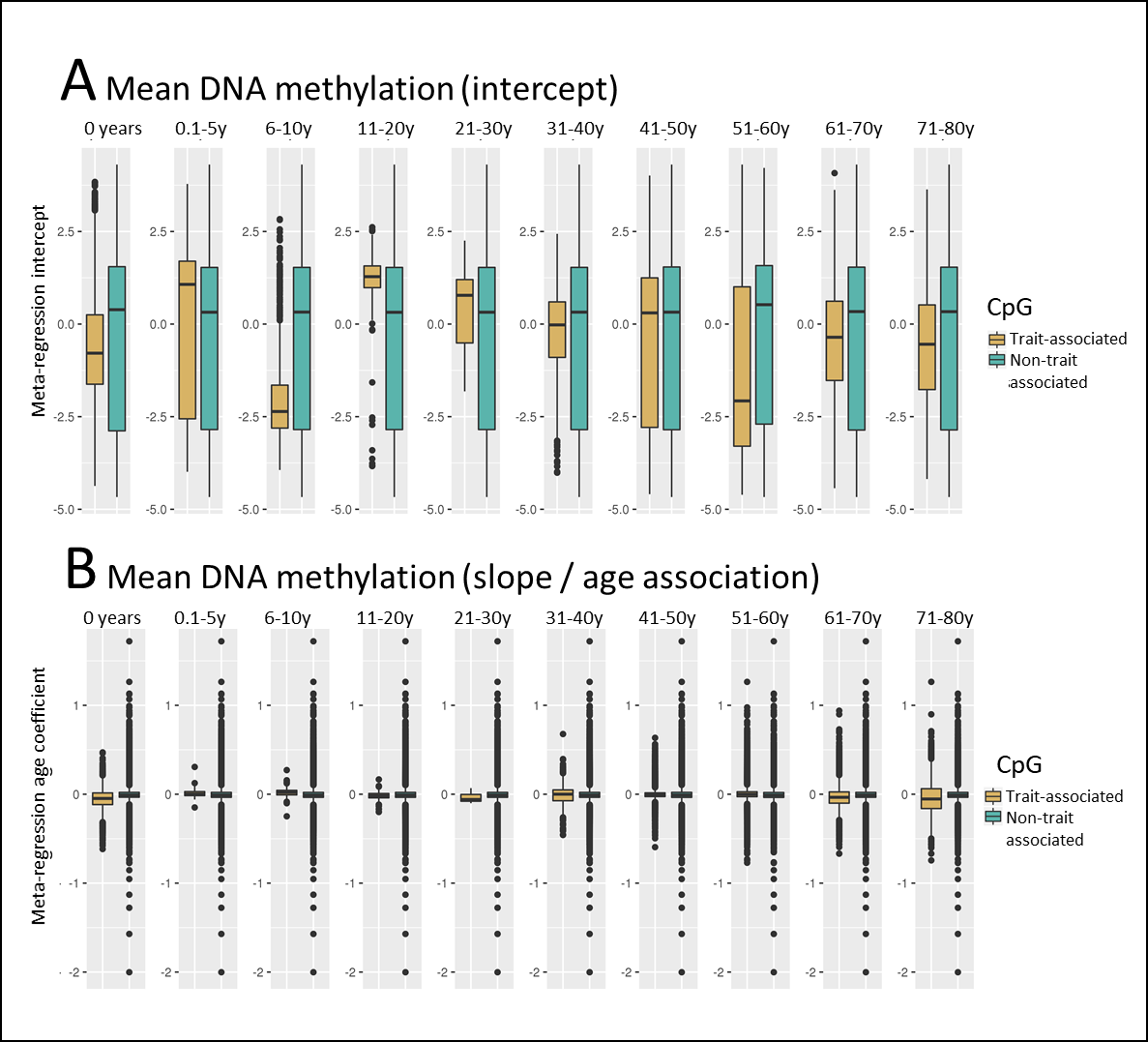

*SM Figure 12*. A) Mean_DNAm_ and B) age~ mean_DNAm_ in CpGs reported in EWAS versus un-reported CpGs by age period (in panels).

**References**

Boyd, A., Golding, J., Macleod, J., Lawlor, D. A., Fraser, A., Henderson, J., Molloy, L., Ness, A., Ring, S., & Davey Smith, G. (2013). Cohort Profile: The ‘Children of the 90s’—the index offspring of the Avon Longitudinal Study of Parents and Children. *International Journal of Epidemiology*, *42*(1), 111–127. https://doi.org/10.1093/ije/dys064

Christiansen, C., Castillo-Fernandez, J. E., Domingo-Relloso, A., Zhao, W., El-Sayed Moustafa, J. S., Tsai, P.-C., Maddock, J., Haack, K., Cole, S. A., Kardia, S. L. R., Molokhia, M., Suderman, M., Power, C., Relton, C., Wong, A., Kuh, D., Goodman, A., Small, K. S., Smith, J. A., … Bell, J. T. (2021). Novel DNA methylation signatures of tobacco smoking with trans-ethnic effects. *Clinical Epigenetics*, *13*, 36. https://doi.org/10.1186/s13148-021-01018-4

Deary, I. J., Gow, A. J., Pattie, A., & Starr, J. M. (2012). Cohort profile: The Lothian Birth Cohorts of 1921 and 1936. *International Journal of Epidemiology*, *41*(6), 1576–1584. https://doi.org/10.1093/ije/dyr197

Elliott, J., & Shepherd, P. (2006). Cohort Profile: 1970 British Birth Cohort (BCS70). *International Journal of Epidemiology*, *35*(4), 836–843. https://doi.org/10.1093/ije/dyl174

Fraser, A., Macdonald-Wallis, C., Tilling, K., Boyd, A., Golding, J., Davey Smith, G., Henderson, J., Macleod, J., Molloy, L., Ness, A., Ring, S., Nelson, S. M., & Lawlor, D. A. (2013). Cohort Profile: The Avon Longitudinal Study of Parents and Children: ALSPAC mothers cohort. *International Journal of Epidemiology*, *42*(1), 97–110. https://doi.org/10.1093/ije/dys066

Harris, P. A., Taylor, R., Thielke, R., Payne, J., Gonzalez, N., & Conde, J. G. (2009). Research electronic data capture (REDCap)—A metadata-driven methodology and workflow process for providing translational research informatics support. *Journal of Biomedical Informatics*, *42*(2), 377–381. https://doi.org/10.1016/j.jbi.2008.08.010

Houseman, E. A., Accomando, W. P., Koestler, D. C., Christensen, B. C., Marsit, C. J., Nelson, H. H., Wiencke, J. K., & Kelsey, K. T. (2012). DNA methylation arrays as surrogate measures of cell mixture distribution. *BMC Bioinformatics*, *13*, 86. https://doi.org/10.1186/1471-2105-13-86

Jones, S., Tillin, T., Park, C., Williams, S., Rapala, A., Al Saikhan, L., Eastwood, S. V., Richards, M., Hughes, A. D., & Chaturvedi, N. (2020). Cohort Profile Update: Southall and Brent Revisited (SABRE) study: a UK population-based comparison of cardiovascular disease and diabetes in people of European, South Asian and African Caribbean heritage. *International Journal of Epidemiology*, *49*(5), 1441–1442e. https://doi.org/10.1093/ije/dyaa135

Maddock, J., Castillo-Fernandez, J., Wong, A., Cooper, R., Richards, M., Ong, K. K., Ploubidis, G. B., Goodman, A., Kuh, D., Bell, J. T., & Hardy, R. (2019). DNA methylation age and physical and cognitive ageing. *The Journals of Gerontology: Series A*. https://doi.org/10.1093/gerona/glz246

Mansell, G., Gorrie-Stone, T. J., Bao, Y., Kumari, M., Schalkwyk, L. S., Mill, J., & Hannon, E. (2019). Guidance for DNA methylation studies: Statistical insights from the Illumina EPIC array. *BMC Genomics*, *20*(1), 366. https://doi.org/10.1186/s12864-019-5761-7

McCartney, D. L., Zhang, F., Hillary, R. F., Zhang, Q., Stevenson, A. J., Walker, R. M., Bermingham, M. L., Boutin, T., Morris, S. W., Campbell, A., Murray, A. D., Whalley, H. C., Porteous, D. J., Hayward, C., Evans, K. L., Chandra, T., Deary, I. J., McIntosh, A. M., Yang, J., … Marioni, R. E. (2019). An epigenome-wide association study of sex-specific chronological ageing. *Genome Medicine*, *12*(1), 1. https://doi.org/10.1186/s13073-019-0693-z

Northstone, K., Lewcock, M., Groom, A., Boyd, A., Macleod, J., Timpson, N., & Wells, N. (2019). The Avon Longitudinal Study of Parents and Children (ALSPAC): An update on the enrolled sample of index children in 2019. *Wellcome Open Research*, *4*, 51. https://doi.org/10.12688/wellcomeopenres.15132.1

Power, C., & Elliott, J. (2006). Cohort profile: 1958 British birth cohort (National Child Development Study). *International Journal of Epidemiology*, *35*(1), 34–41. https://doi.org/10.1093/ije/dyi183

Reinius, L. E., Acevedo, N., Joerink, M., Pershagen, G., Dahlén, S.-E., Greco, D., Söderhäll, C., Scheynius, A., & Kere, J. (2012). Differential DNA methylation in purified human blood cells: Implications for cell lineage and studies on disease susceptibility. *PloS One*, *7*(7), e41361. https://doi.org/10.1371/journal.pone.0041361

Relton, C. L., Gaunt, T., McArdle, W., Ho, K., Duggirala, A., Shihab, H., Woodward, G., Lyttleton, O., Evans, D. M., Reik, W., Paul, Y.-L., Ficz, G., Ozanne, S. E., Wipat, A., Flanagan, K., Lister, A., Heijmans, B. T., Ring, S. M., & Davey Smith, G. (2015). Data Resource Profile: Accessible Resource for Integrated Epigenomic Studies (ARIES). *International Journal of Epidemiology*, *44*(4), 1181–1190. https://doi.org/10.1093/ije/dyv072

Shah, S., McRae, A. F., Marioni, R. E., Harris, S. E., Gibson, J., Henders, A. K., Redmond, P., Cox, S. R., Pattie, A., Corley, J., Murphy, L., Martin, N. G., Montgomery, G. W., Starr, J. M., Wray, N. R., Deary, I. J., & Visscher, P. M. (2014). Genetic and environmental exposures constrain epigenetic drift over the human life course. *Genome Research*, *24*(11), 1725–1733. https://doi.org/10.1101/gr.176933.114

Sugden, K., Hannon, E. J., Arseneault, L., Belsky, D. W., Corcoran, D. L., Fisher, H. L., Houts, R. M., Kandaswamy, R., Moffitt, T. E., & Poulton, R. (2020). Patterns of Reliability: Assessing the Reproducibility and Integrity of DNA Methylation Measurement. *Patterns*, 100014.

Taylor, A. M., Pattie, A., & Deary, I. J. (2018). Cohort Profile Update: The Lothian Birth Cohorts of 1921 and 1936. *International Journal of Epidemiology*, *47*(4), 1042–1042r. https://doi.org/10.1093/ije/dyy022

Tillin, T., Forouhi, N. G., McKeigue, P. M., Chaturvedi, N., & SABRE Study Group. (2012). Southall And Brent REvisited: Cohort profile of SABRE, a UK population-based comparison of cardiovascular disease and diabetes in people of European, Indian Asian and African Caribbean origins. *International Journal of Epidemiology*, *41*(1), 33–42. https://doi.org/10.1093/ije/dyq175

Verdi, S., Abbasian, G., Bowyer, R. C. E., Lachance, G., Yarand, D., Christofidou, P., Mangino, M., Menni, C., Bell, J. T., Falchi, M., Small, K. S., Williams, F. M. K., Hammond, C. J., Hart, D. J., Spector, T. D., & Steves, C. J. (2019). TwinsUK: The UK Adult Twin Registry Update. *Twin Research and Human Genetics: The Official Journal of the International Society for Twin Studies*, *22*(6), 523–529. https://doi.org/10.1017/thg.2019.65

Viechtbauer, W. (2010). Conducting Meta-Analyses in *R* with the **metafor** Package. *Journal of Statistical Software*, *36*(3). https://doi.org/10.18637/jss.v036.i03

Zhang, Q., Marioni, R. E., Robinson, M. R., Higham, J., Sproul, D., Wray, N. R., Deary, I. J., McRae, A. F., & Visscher, P. M. (2018). Genotype effects contribute to variation in longitudinal methylome patterns in older people. *Genome Medicine*, *10*(1), 75. https://doi.org/10.1186/s13073-018-0585-7
